# How the brain negotiates divergent executive processing demands: Evidence of network reorganization during fleeting brain states

**DOI:** 10.1101/2020.05.30.125476

**Authors:** Mengting Liu, Robert A. Backer, Rachel C. Amey, Chad E. Forbes

## Abstract

During performance in everyday contexts, multiple networks draw from shared executive resources to maintain attention, regulate arousal, and problem solve. At times, requirements for attention and self-regulation appear to be in competition for a “limited pool” of resources. How does the brain attempt to resolve conflicts arising from multiple processing demands? In the present study, participants were exposed to either a stress or control prime, after which electroencephalographic (EEG) activity was recorded as they solved math problems. Phase-locking was examined within four networks implicated in math-solving and evaluative stress: frontopareital (FP), default mode (DM), emotion generation (EG), and emotion regulation (ER) networks. Findings revealed differing strategies, depending on the presence of stress: states dominated by frontopareital and emotion regulation network dynamics supported optimum performance generally, while during stress, states dominated by emotion regulation and default mode networks are more important for performance. Implications for networks’ cooperative dynamics and DMN’s role in coping are considered.

## Introduction

Many tasks require negotiating processing demands that appear to be at odds with one another, yet how the brain manages to accomplish this remains somewhat of a mystery. One example is problem solving, which requires both efficiency in cognitive processes like attention and working memory, as well as regulating arousal, which often increases with task difficulty or evaluative threats—both of which rely on common executive processing resources. How does the brain manage these overlapping demands simultaneously, and what characterizes better performance under such conditions? To clarify these relationships, the present study contrasted the frontopareital network (FPN)—important for performing calculations—with additional networks involved in self-referent processing (DMN) and emotion regulation (ERN) as subjects solved math problems in the presence or absence of stressors. Hidden Markov Modeling (HMM) was applied to encephalography (EEG) data at the network level (network synchrony values) to identify brain states that differed based on these networks’ levels of activity, thus providing a functionally relevant measure of state composition. Findings suggested two routes to success as problem solving difficulty increased: FPN and ER dominant states were important generally, but for those under stress, states marked by high ER and DMN predicted optimal performance. Taken together, these findings suggest that, while problem-solving itself requires canonical FPN efficiency, individuals who perform better under stress are *also* able to recruit DMN in order to negotiate additional intrusions that would otherwise undermine their work.

### Competition for Executive Resources During Problem Solving

Neuroscience literature offers multiple examples of situations where two networks appear to be locked in competition with each other and must achieve a balance. For example, we know that basal ganglia and hippocampal systems compete, and that the FPN and DMN are frequently associated with a contest for internally vs externally oriented attention. In stressful performance contexts, there appears to be a similar relationship between managing arousal (associated with ERN) and orienting oneself to the “task at hand” (reliably linked to FPN). Since both networks draw from common executive resources in dorsolateral prefrontal cortex (DLPFC), it stands to reason that as FPN and ERN are engaged at the same time, people perform poorly. Conventionally, this has been explained as resulting from a “tug-of-war” for limited resources (e.g., Fig 1), and indeed, research has shown that co-activation of or cooperation between FPN and ERN often attenuates performance as difficulty and arousal increase.

**Figure 1.**
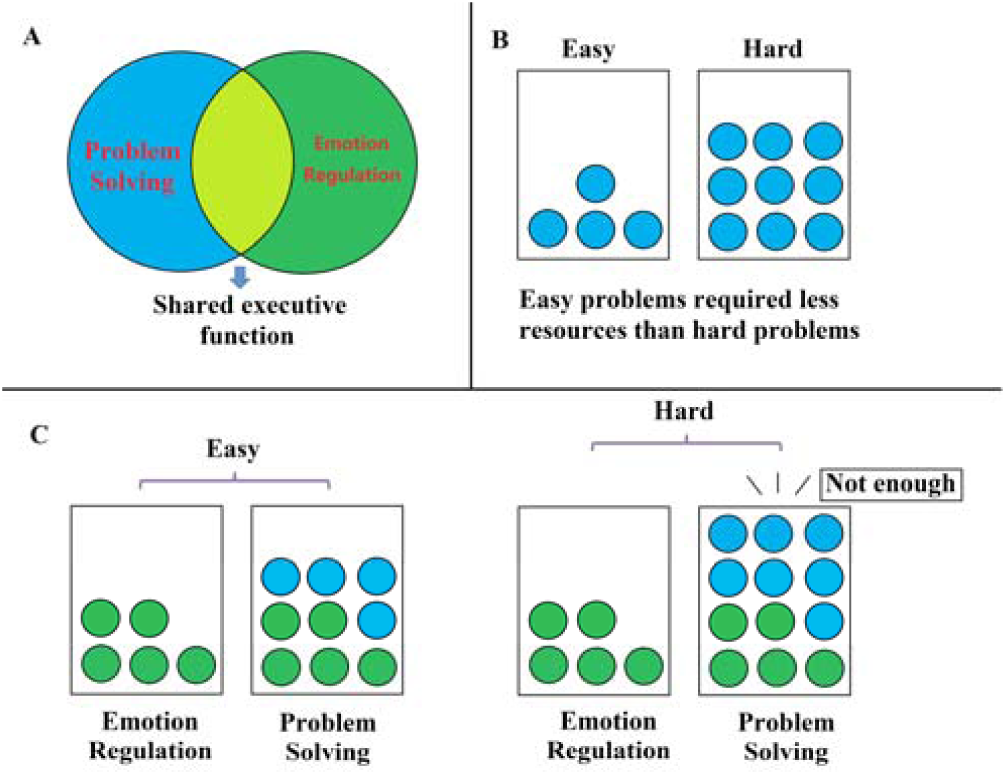
One of the hypothesis for the relationship between emotion regulation and problem solving -- competiting for limited executive resources.

Examining further, problem solving relies on coordination between visual, motor, and higher-order cognitive processes. For example, Anderson, Lee, & Fincham (2014) have demonstrated how solving algebra problems occurs through a collection of functionally relevant states (*define, encode, transform, compute*, and *respond*), each drawing in differing proportions, from—among others—visual attention, executive control, metacognitive, and motor network regions. Better performance was predicted to the extent that states were dominated by essential areas, which are reliably linked to various aspects of math solving. Moreover, networks also must be able to dynamically reconfigure, in order to move from one state of processing to the next, sometimes iteratively (Anderson, Lee, & Fincham, 2014). This underscores that “optimum ingredients” for performance are time-locked. That is, what is helpful at one stage of processing (i.e. one state) may be counterproductive if “leaking” into other stages. Indeed, in the case of arousal or stress during performance, ill-timed activation of emotion or self-related brain regions during problem solving is precisely what is conceptualized to undermine performance (van Ast et al., 2016; Amey et al., 2018). This is also reflected in work associating stress during performance with too many or too few network states, to the decrement of performance outcomes (Liu, Amey, Forbes, 2017; Anderson et al., 2011).

Yet, while prior work has characterized what is *bad* for performance on it’s own, we have yet to explore what might actually be *good* under realistic conditions where individuals must contend with everyday stressors during problem solving. Certainly, some individuals fare better than others in stressful performance, presumably due to network dynamics that make it possible to manage conflicting executive demands. Borrowing from negotiation terminology, it is possible that instead of the *zero-sum* situation articulated thus far, additional dynamics may emerge to effectively *“widen the pie”* such that disparate networks are better able to cooperate. Many paradigms do not deliberately manipulate stressors with the goal of searching for predictors of better outcomes, and thus may fail to unearth such processes. In the following section, we review performance under stress, and discuss potential palliative mechanisms at play.

### Possible Modes of Network Cooperation During Problem Solving under Stress

In stressful performance contexts, several abilities are often important: focus for solving actual problems—as has thusfar been discussed—as well as regulating arousal (e.g., the canonical Yerkes-Dodson law, 1908), and managing evaluative threats (Liu et al., 2020). There is ample evidence that when confronted with evaluative threats, people have intrusive self-doubts (Cadinu, Maass, Rosabianca, & Kiesner, 2005), which may initiate a cycle of hyper-vigilance for critical feedback and exacerbate negative arousal, further detracting from performance (Schmader, Johns, & Forbes, 2008). When confronted by self-doubts, two kinds of self-reflective responses may ensue: one may begin to self-denigrate (e.g. “I’m messing up again”, “I do this all the time”, “I’m not good at this”) or self-affirmation (e.g. “I have done well on other things like this”, “I have the ability to handle this”). Thus, in addition to the activities of problem solving and managing stress, it may also be important to have an autobiographical memory system capable of retrieving positive self-content to assist in mitigating evaluative threats.

One candidate for such a role is the DMN, which is active during internally oriented processing like autobiographical memory, and, in stressful situations, may serve a coping role by enabling people to recall positive memories as a buffer to identity threats (Spreng, Mar & Kim 2011). Internally oriented processing is often thought of as reflecting on one’s past actions, contemplating social relationships, and sometimes mind wandering. Thus, DMN has often been thought to conflict with externally focused attention and FPN-linked problem-solving. However, work has shown that DMN and FPN are not always anti-correlated, but can even work together during certain mental calculations like autobiographical memory tasks. Moreover, greater resting-state DMN synchrony has also been found to predict accuracy on post-task self-appraisals when individuals were exposed to evaluative threats during performance, further suggesting a role in coping. Self-awareness and coping are not typically considered regarding performance, but certainly plan an online role when significant stressors are present. Thus we posit that, for better performance in stressful contexts, three cognitive components are likely relevant, which we loosely relate to the 3 networks discussed thus far: executive attention (FPN), emotion regulation (ERN), and a self concept, capable of coping or putting evaluative threats into perspective (possibly DMN).

### Further Examining Network Dynamics during Stressful Performance

Previous work on optimum ingredients for performance conventionally tell us that more FPN is good, while less ERN and DMN are counterproductive. Moreover, work on stress indicates that frontopareital integrity decreases, while emotion regulation areas instead come to predominate (Liston, McEwen, & Casey, 2009). However, we argue that defining the relationships in this ratio may not be giving us the full picture, since when ERN or DMN are couched as “at the expense of” FPN and viewed in experimentally neutral contexts, results are more likely to capture distraction. Instead, by exploring the roles of these networks in more ecologically-valid situations that place their related executive processes in direct conflict (i.e. performance where stress is high), we may gain a somewhat different story. Moreover, prior functional magnetic resonance imaging (fMRI) work has largely looked at neural activity over large time blocks. Under such conditions, it is more likely that we would find the conventional pattern in Fig. 2A, yet neural states can be fleeting, and therefore it is possible that positive roles for ERN or DMN would be detected on shorter time scales, such as those measureable with EEG. The purpose of the present study is to examine what network dynamics emerge in this circumstance.

**Figure 2.**
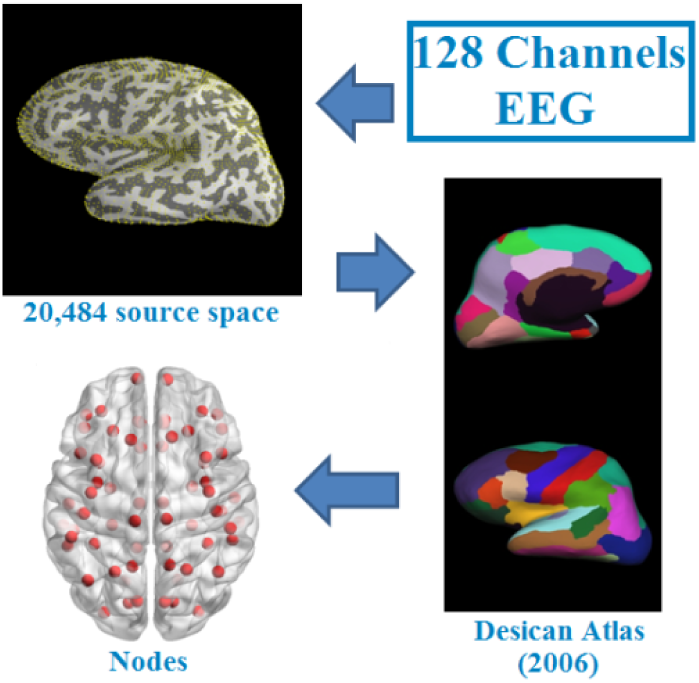
Construction of adjacency matrix cortical source space

Rather than individual regions, we focused on activation at the level of networks, which can enable us to characterize relationships between the broader cognitive processes we are interested in. Based on prior work, we anticipated that FPN would be required for problem-focused deliberation. We hypothesized that some nuanced interaction between networks would emerge as problem difficulty increased (relating to greater arousal), and in the presence of evaluative stressors. Because the “task at hand”, problem solving, would be largely reflected by FPN activity, it was necessary to adopt an approach capable of identifying fleeting brain states that might reflect more nuanced relationships between networks. We thus employed Hidden Markov Modeling (HMM) to individual-level data, capable of distinguishing states that predict better performance in the presence or absence of stressors (for reference, see Liu, Amey, & Forbes, 2017, and Anderson, Lee, & Fincham, 2014). We considered two possible dynamics that might emerge in states: states might either be dominated by a particular network (reflecting a particular functional process) or reflect greater cooperation *between* two networks.

Specifically, we manipulated identity threat, a robust stressor closely tied to evaluative threats (Schmader, Johns, Forbes, 2008), across math problem solving experimental conditions. In math, negative stereotypes about women’s’ abilities prime threat cues, inducing a maladaptive cycle of hypervigilance and attempts to regulate accompanying emotions. Past work (Liu, Amey, & Forbes, 2017) demonstrated that this paradigm is capable of inducing negative arousal in association with a quadratic trend in neural state entropy and poorer performance. As such, identity threat provides a realistic, evaluative threat stimulus strongly tied to self-related cognition. Thus, women across experimental cells would not differ in any other way, and men provide a contrast that allows for confirmation that identity indeed drove any effects. Men and women completed math problems while EEG activity was recorded, affording maximum temporal sensitivity. This study was the first we know of to examine neural states in short EEG timescales, while pitting processing demands against one another to assay what promotes better performance. As such, the nature of our HMM approach is best described as exploratory and data-driven. However, past literature did permit us to make some tentative guesses about what kinds of functional themes might emerge. We expected to find that FPN would predict performance on easier problems, in line with canonical expectations for math solving. As difficulty increased, we considered that either FPN alone, or in concert with ERN—to manage greater accompanying arousal—would continue be important for the general population (men and non-stressed women). Finally, we hypothesized that, for women (and not men) in the stress condition, individual differences in states characterized by some nuanced relation in FP, ER, or DM networks would predict better performance, reflecting efficiency in managing evaluative threats.

## Methods

### Participants

One hundred and fifty-seven white participants (84 females) completed this study for payment. We recruited only participants who were aware of a negative female math stereotype. Specifically, participants needed to score a three or lower on the following question during a pre-study screening in order to qualify for the current study: “Regardless of what you think, what is the stereotype that people have about women and men’s math ability” (1= Men are better than women; 7= Women are better than men).” Four participants were excluded because their EEG data lacked more than half of the math solving trials. One further participant was excluded for having most of the math solving trials shorter than three seconds, likely to reflect arbitrary answers, rather than engagement in actual math solving. Finally, 152 participants (82 females) were remained for our analysis.

### Procedure

Upon entering the experiment room, participants were taken to a soundproofed EEG chamber, seated in front of a computer, and prepared for electroencephalographic recording. Following a previously established SBS manipulation (Forbes et al., 2019), participants were randomly assigned to either a “diagnostic math test” condition (DMT; SBS condition) or a “problem solving” condition (PST; control condition). In the DMT condition, participants were informed that results from the following tasks would be diagnostic of their math ability. In the PST condition, participants were informed that results would be diagnostic of the types of problem-solving techniques they prefer. To further prime stereotype threat, participants marked their gender in the DMT condition, all DMT sessions had a male experimenter present, and all instructions were read to them via a male experimenter’s voice (Forbes et al., 2018). Conversely, participants in the PST condition did not mark their gender, all PST sessions had all female experimenters present and all instructions were read to them via a female experimenters voice. Following the instructions, participants completed a math feedback task for 34 minutes. Participants then answered a series of post-task questionnaires, after which they were debriefed and paid for their participation.

### Math Feedback Task

Participants completed a 34-minute math task identical to Forbes, Amey, Magerman, Duran, & Liu (2018). The task consisted of standard multiplication and division problems (e.g. 7×20=) presented on the computer screen. In a given trial, participants were provided with three answer options for each problem (A, B, or C). The correct answer to each problem was placed randomly in one of the three answer positions. Participants answered each multiple choice problem using a button box placed on their laps and were not permitted to use scratch paper. After answering, participants received feedback on the screen for 2 seconds (“Correct” or “Wrong”). After feedback, the next problem was presented. Each problem was displayed for a maximum of 17 seconds. If participants did not answer a problem within this period, they would, by default, receive negative feedback (“Wrong”). On average, participants completed 83.9 problems. Our measure of math score accuracy was calculated by dividing the total number of correct responses by the total number of attempted problems and multiplying that outcome by 100.

### EEG Recording

Continuous EEG activity was recorded using an ActiveTwo head cap and the ActiveTwo Biosemi system (BioSemi, Amsterdam, Netherlands). Recordings were collected from 128 Ag-AgCl scalp electrodes and from bilateral mastoids. Two electrodes were placed next to each other 1 cm below the right eye to record startle eye-blink responses (Liu et al., 2020; Liu et al., 2017). A ground electrode was established by BioSemi’s common Mode Sense active electrode and Driven Right Leg passive electrode. EEG activity was digitized with ActiView software (BioSemi) and sampled at 2048 Hz. Data was downsampled post-acquisition and analyzed at 512 Hz. EEG signals were epoched and stimulus locked to 500 ms before participants were presented with a math problem to the time they solved the math problem. EEG artifacts were removed via FASTER (Fully Automated Statistical Thresholding for EEG artifact Rejection) (Nolan, Whelan, & Reilly, 2010), an automated approach to cleaning EEG data that is based on independent component analysis (ICA) and incorporates multiple statistical steps. Specifically, raw EEG data were initially filtered through a band-pass, finite impulse response (FIR) filter between 0.3 and 55 Hz. First, EEG channels with significantly unusual variance (operationalized as activity with an absolute z score larger than 3 standard deviations from average), average correlation with all other channels, and Hurst exponent were removed and interpolated from neighboring electrodes using spherical spline interpolation function. Second, EEG signals were epoched and baseline corrected, and epochs with significant unusual amplitude range, variance, and channel deviation were removed. Third, the remaining epochs were transformed through ICA. Independent components with significant unusual correlations with EMG channels, spatial kurtosis, slope in filter band, Hurst exponent, and median gradient were subtracted and the EEG signal was reconstructed using the remaining independent components. Finally, EEG channels within single epochs that displayed significant unusual variance, median gradient, amplitude range, and channel deviation were removed and interpolated from neighboring electrodes within those same epochs.

### Source Localization

In order to map electrodes signals to activity in neural regions, forward model and inverse models were calculated with MNE-python, an open access software (Gramfort et al., 2013 and Gramfort et al., 2014). Forward model solutions for all source locations located on the cortical sheet were computed using a 3-layered boundary element model (BEM) (Hämäläinen and Sarvas, 1989), constrained by the default average template of the anatomical MNI MRI. Cortical surfaces extracted with FreeSurfer were sub-sampled to about 10,242 equally spaced vertices on each hemisphere. Further, a noise covariance matrix for each individual was estimated from the pre-stimulus EEG recordings after the preprocessing. Forward solution, noise covariance and source covariance matrices were used to calculate the dynamic statistical parametric mapping (dSPM) estimated (Dale et al., 2000) inverse operator (Dale et al., 1999). Inverse computation was done using a loose orientation constraint (loose = 0.11, depth = 0.8) (Lin et al., 2006). Surface was divided into 68 anatomical regions of interest (ROIs; 34 in each hemisphere) based on the Desikan–Killiany atlas (Desikan et al., 2006). For each participant, a time course was calculated for each area/node by averaging the localized EEG signal of all of its constituent voxels at each time point during task performance.

### Frequency Selection

Managing cognitively demanding tasks in stressful situations depends upon a number of cognitive resources: cognitive control, working memory, emotion regulation, attention, etc. Such functions are supported by contributions from multiple neural regions, which operate within networks. Network activity is diversely reflected through regions’ *coherence*. That is, regions communicate through neural synchrony, and as groups of neurons become entrained for a given purpose, their oscillatory firing rates align at various frequencies.

For instance, fronto-parietal coherence in theta (4–8 Hz) and upper alpha (8-12 Hz) frequency ranges has been found to reflect working memory functioning (Sauseng et al., 2005). Additionally, past work also demonstrates converging evidence that working memory persistence is supported by coherence in the theta and gamma (30–100 Hz) ranges (Lisman, 2010; Raghavachari et al., 2001; Liu et al., 2017) in frontal and parietal cortex.

A number of emotion-related processes are also reflected in oscillatory activity. For instance, amygdala neurons display elevated theta band activity during emotional arousal (Denis Paré et al., 2002). Emotion processing—especially negative arousal processes—was long thought to be primarily instantiated by alpha band amygdala activity (Aftanas et al., 2005). However, it has been recently identified that long-range interaction of prefrontal structures and the amygdala occurs within in the theta. Moreover, past research found an increase in frontal theta oscillations during emotion regulation and a relationship between frontal theta power and the subjective success of emotion regulation (Ertl et al., 2013). Across the frontal cortex, theta activity, appears to reflect a common medium for carrying out cognitive control (Cavanagh & Frank, 2014).

Given the multiplicity of cognitive components involved in performance under SBS, it was necessary that our frequency band selection allowed for the examination of diverse neural functions. In contrast to other bands, theta activity has been consistently incorporated across different components relevant to math solving within emotion contexts. Hence, we chose to examine neural activity within the theta band in this study, which allows us to examine the relationships of these functions on a common frequency scale.

### Time varying functional connectivity estimation and network construction

EEG data was collected from 150 participants (79 threat, 71 control). Each math problem—one trial—was presented for a maximum of 17 seconds, during which participants could provide their answer. EEG data was only analyzed within the time period they actually solved problems (i.e. until they gave their answer). Trials with solving times of less than 3 seconds were excluded, because these trials were likely to have reflected arbitrary answering rather than serious solving of math problems. After pruning short trials, 10,950 math-solving trials remained.

Separate time series were extracted for all 68 sources on each trial, using MNE. For all pairings of the 68 sources, dynamic functional connectivity between a given pair of two sources was computed across the time series by using a sliding window approach (Hutchison et al., 2013). Time series were divided into 128 smaller temporal points, and a window of 1 second in length was applied in steps (0.0625 seconds each) across the series from beginning to end. The two band-power spectra of each source pair, within a given time window, were cross-correlated using Pearson’s correlation coefficient (Chen, Ros & Gruzelier, 2012). In other words, for each math-solving trial, we obtained a symmetric 68 × 68 connectivity score matrix for each of the time window periods. Each matrix was first converted to a sparser, positive-only matrix by removing all negatively weighted edges (i.e. pairs of connections between sources) due to the ambiguous meaning of negative connectivity (Rudie et al., 2013). The remaining matrix was further pruned by applying a statistical threshold (Liu et al., 2011, 2013) to retain coefficients *r*_*ij*_ <= 50% of the total positive connections. Specifically, we further optimize our model by examining activity over several smaller functional *subnetworks* (subsets of the entire matrix) relevant to understanding performance under stress.

### Functional sub-networks

Problem solving under stress requires a number of cognitive resources, e.g. executive function (EF), working memory (WM), attention, emotion, emotion regulation and self-regulation. We selected the fronto-parietal network (FPN) as a proxy for EF, WM, and attention.

Emotion consists of a set of reactions triggered by affective internal or external stimuli (Gross, 1998; Kohn et al., 2014), which may be separated into two major components, namely emotion generation and emotion regulation (Kohn et al., 2014; Goldin et al., 2008; Gross & Barrett, 2011). it’s long been proposed that emotion generation engages the limbic system (LeDoux, 2000; Phan et al., 2003), such as amygdala and ventral striatum. Recent studies also found some closely related cortical regions such as anterior insula and anterior cingulate cortex (ACC; Etkin et al., 2015). Particularly, identity threat stimulated emotion may start from an independent type of emotion called top-down emotion generation (Ochsner et al., 2009), which is closely associated with medial prefrontal cortex (mPFC)/ventral ACC (vACC) (Silvers et al., 2015). Due to the limited spatial resolution for reconstructing brain activity in deep sources like amygdala in EEG techniques, we only selected insula and vACC and created an emotion generation (EG) network. On the other hand, emotion regulation refers to a conscious or unconscious process to evaluate, modulate or alternate emotion trajectory (Etkin, Buchel & Gross, 2015). Meta analysis of neuroimaging studies has revealed that a number of brain regions, including dorsolateral prefrontal cortex (DLPFC), ventrolateral PFC (VLPFC), vACC, dorsal ACC (dACC), mPFC, orbital PFC, and precuneus (especially for self-regulation) (Wager et al, 2008;Ochsner & Gross, 2005; Buhle et al, 2014; Phillips, Ladouceur & Drevets, 2008; Ochsner et al., 2004;Kohn et al., 2014; Ochsner et al., 2002; Ferri et al., 2015). Hence, we engage these regions to our emotion regulation network (ER).

Furthermore, default mode network (DMN) was also added to our analysis, not only for its role of task-negative network—which can be considered as a control for EF network—but also for the recent findings of its character in emotion coping and self-regulation(Forbes et al., 2015; Jordano & Touron 2017; Ossandon et al., 2011; Leitner & Forbes, 2015).

All brain regions engaged in four sub-networks were selected from the one of the 68 whole-brain atlas. Figure 3 represents all the brain regions involved in each network.

**Figure 3.**
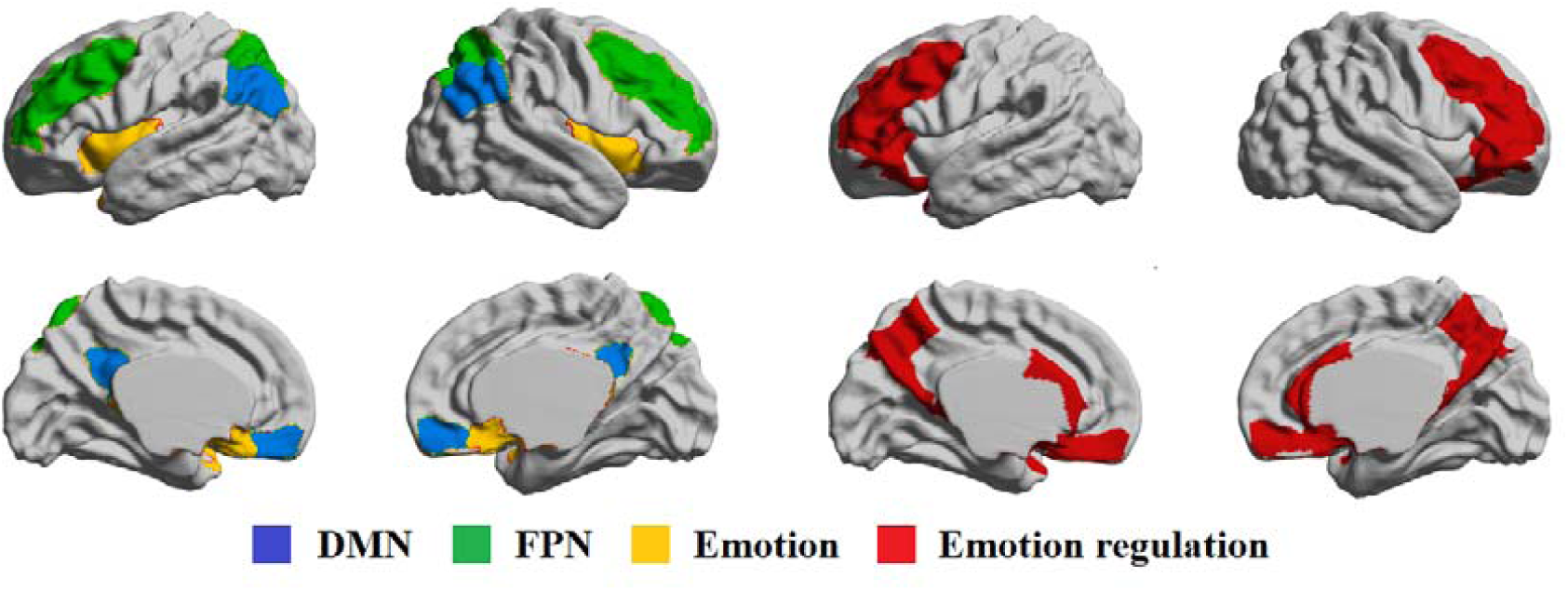
Brain regions involved in four sub-networks

### Graph measure for each subnetwork

The brain is a complex system, it consists of several functional *modules*, or nearly decomposable units (Simon, 1991). This structure aggregates subsystems that can perform independent functions without interacting the rest of the system (Bessett, et al., 2011). In other words, modules are groups of brain regions with many intra-module links, but few inter-modular links to external groups. It has been found that brain networks typically show high modular architectures. In brain networks, topological modules are often made up of anatomically neighboring and/or functionally related cortical regions, inter-module connections tend to appear in relatively long distance (Meunier, Lambiotte & Bullmore, 2010). Brain regions or the constituent nodes of topological modules are often anatomically co-localized in the brain (Bertoleroa, Yeo & D’Espositoa, 2015). This arrangement seems to be advantageous in terms of minimizing the connection distance or wiring cost of intra-modular edges. For example, modularity analysis conducted on whole brain network will always yield fronto-temporal, central, parietal, occipital and default-mode modules at the highest level of the hierarchy. Little is known however, that how the topological modularity of large-scale brain networks is related to other aspects of modularity psychologically (Meunier, Lambiotte & Bullmore, 2010). For example, a brain network that is psychologically/cognitively meaningful (e.g. working memory network) can be widely distributed among several anatomical modules. Traditional modularity analyses cannot answer the question of whether regions in psychologically meaningful network would work together in a more efficient and close manner during cognitive tasks. To address this question of modular organization within psychological communities, we apply a novel measure of modularity that extends the rationale of modularity in whole network analyses to a priori defined *sub*-networks, thought to play integral roles in psychological processes of interest. We define this new measure as *selected modularity* (Forbes et al., 2018).

Selected modularity is a measure that compares within module connectivity (within a predefined sub-network) to between module connectivity (operationalized as Pearson correlation coefficient between pre-defined subnetworks or other regions across the whole brain network). Selected modularity was calculated using the equation below:

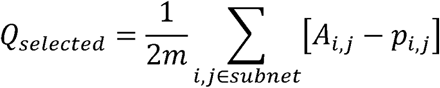

Selected modularity measures the integrated level or within network connectivity of a subnetwork of interest with respect to either another subnetworks of interest, or the whole brain (in our case it’s with respect to the whole brain). Higher selected modularity indicates that the nodes within a subnetwork are more efficiently interconnected with one another, compared to other nodes in the whole brain as well as connections that would be expected by chance. Therefore, higher selected modularity suggests that a network was more active or efficient during a given cognitive process; obtaining this measure at each temporal point affords us an index of networks’ dynamic roles during cognition.

### Brain States Identification

After the estimation of whole-brain dynamic functional connectivity and dynamic graph analyses, the obtained dynamic graph patterns were summarized into a smaller set of graph states, which allows identification of recurring spatio-temporal graph patterns (Preti, Bolton & Ville, 2017). Several machine-learning methods have been demonstrated for discovering neural states, such as k-mean clustering (Allen et al., 2012; Mantena et al., 2009), hierarchical clustering (Ou et al., 2013), or modularity approaches (Yu et al., 2015). In this study, we utilized Hidden Markov Model (HMM) to identify these states, as it confers unique advantages over other methods (e.g. Chen, et al., 2016).

### *Hidden Markov Model (HMM).* (new model used)

An HMM is a state-space stochastic model. Let s_t_ and o_t_ denote assignment of a hidden state and an observation at time t, respectively. Hidden and observable processes over T time-length can be denoted by {s_l_,·, s_t_,·, s_T_} and {o_l_,·, o_t_,·, o_T_}. For the hidden process, we model it with a first-order Markov chain, which means the hidden state at time t, i.e., s_t_, is dependent on the hidden state at time (t − 1), i.e., s_t−l_. The observable variable at time t, i.e., o_t_, is dependent on the hidden state at the same time, i.e., s_t_.

Formally, when we consider a number *K* of hidden states, a hidden process is defined by two probability distributions: state transition probability and initial state probability. A state transition probability A= [a_ij_]_i,j∈{l,…,K}_, where a_ij_ = P(s_t_ = j|s_t−l_ = i), denotes the probability of changing from one hidden state (s_t−l_ = i) to another hidden state (s_t_ = j). When i = j, suggesting the state doesn’t transition, the state transition probability is replaced by a duration probability D= [P_i_(t − τ)] for a specific hidden state, where τ is the initial time for the current specific hidden state, and t− τ represents the duration of the current state. In this work, we use a Gaussian distribution for duration probability. Also an initial state probability Π = [π_i_]_i∈{l,…,K}_, where π_i_ = P(s_l_ = i), which models the probability of starting from a specific hidden state at time t = 1, denotes where the state transition begins. The observable process is depicted by an emission probability density function B = [b_i_]_i∈{l,…,K}_, where b_i_ = P(o_t_ = x_t_|s_t_ = i), which denotes the likelihood of observing the specific observation x_t_ when residing at the hidden state of i. We also use a single Gaussian distribution for emission probability. Thus, an HMM is defined by the parameter set of λ = (Π, A,B, D) (Rabiner, 1989; Suk, et al., 2016).

### Determination of HMM states

HMM was trained using the data from all 10,950 math solving trials. The number of states in this study were determined via a threshold for marginal increase of log-likelihood for the whole dataset (Celeux, 2007). That is, the number of state N was selected when the log-likelihood increased less than 1% with respect to the number of state N+1. Based on this criteria, we selected 10 as the optimal number of states to construct the model. In order to avoid the errors in determining the number of states caused by different methods, and also to avoid the random effect that appears when only 10 states are used, we extend our analyses to 8 to 12 states. Patterns were confirmed only when they were repeatedly found within all these numbers of states.

To first construct HMM models, a learning phase was applied, where an individual HMM model was optimized for best fit of the data. This model was trained via an efficient method proposed by Yu and colleagues (Yu & Kobayashi, 2003, 2006). HMM analyses were conducted on time periods during which individuals solved math problems using the evoked selected modularity curve elicited from the 4 cortical networks outlined in our source model. Once the HMM models were constructed, a second decoding was applied to the selected modularity curves stemming from each source in order to identify what is the most likely state sequence in the model that produced the observations. We obtained a series of states generated from the selected modularity curve in every trial. Figure 4 illustrates the flow chart of HMM analysis in our study.

**Figure 4.**
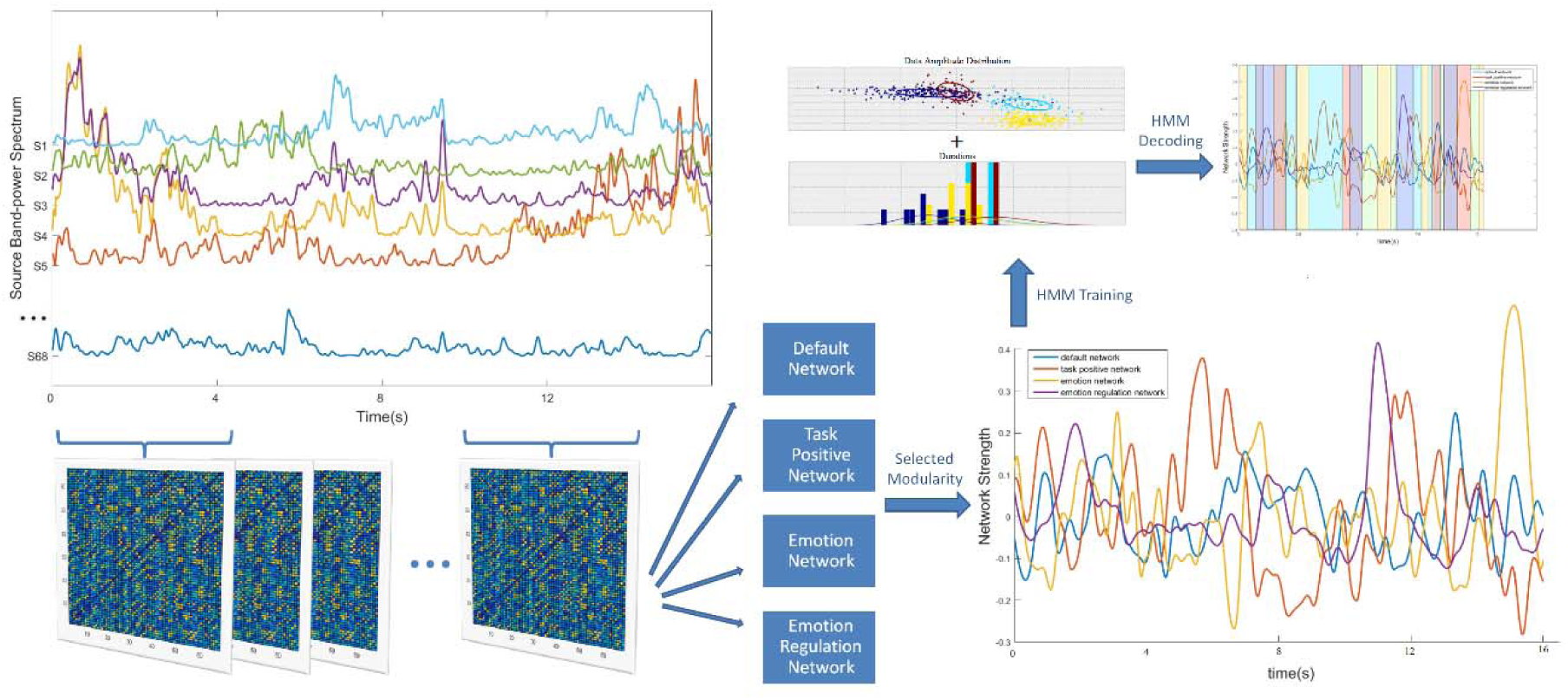
Flowchart of the pipeline for HMM analysis.

Drawing upon past research on dynamic neural states (Hutchison & Morton, 2015), the following three individual measurements assessed state expression separately for each participant during task: (1) duration occupancy (DO), measured as the proportion of all windows labeled as instances of particular states, and computed separately for each of the states; (2) mean dwell time occupancy (MDTO), measured as the proportion of average number of consecutive windows labeled as instances of the same state, and computed separately for each of the states; and (3) the number of states/transitions (NS), measured as the number of states across all math solving tasks.

### Brain state measures

Together with the state distributions, the HMM inference estimates the time courses of the visits to each of the brain states. We used these to look at the extent to which the temporal characteristics of different cognitive states differed in each group. The three features used to characterize the brain states are: the fractional occupancies (reflecting the proportion of time spent in each state); the mean lifetimes (or dwell times, i.e. the amount of time spent in a state before moving into a new state); and the interval times between consecutive visits to a state.

## Results

### Stress manipulation check

Initial analyses on amygdala activity (assessed via startle probes elicited to positive and negative feedback and operationalized as a measure of stress) were conducted in Forbes et al. (2018). These analyses indicated that all participants elicited marginally greater amygdala responses to negative feedback received on the math feedback task compared to positive feedback and women elicited larger amygdala responses to feedback compared to men. However, only women under the stress condition exhibited a unique non-linear (quadratic) relationship in their amygdala responses to feedback over time, suggesting a unique stress response among individuals under conditions of evaluative threat. Women under stress also performed worse on the math task compared to all other conditions (planned contrast on math test accuracy: t (1, 156)=3.17, p=.002, d=.51). These behavioral results, overall, provide supporting evidence that the evaluative threat and thus stress induction manipulation was successful (see details in supplementary results).

### Ten states were identified using HMM

Using EEG trials math solving data from 152 subjects, mapped to four subnetwork based on a 68-region parcellation using dSPM estimated inverse operator, we identified 10 HMM states using Bayesian Hierarchical Hidden Markov Models (HMM). Essentially, this technique finds, in a completely data-driven way, recurrent patterns of network (or HMM state) activity. Each HMM state has parameters describing brain activity in terms of network activity covariations. The method provides information that is temporally resolved (different networks are described as being active or inactive at different points in time). Importantly, while the spatial and temporal description of the states is common to all subjects, each subject has their own state time course, representing the probability of each HMM state being active at each instant. See Fig. 3 for a graphical example, and an illustration of the entire pipeline.

### Transition probabilities of brain states

Next, we investigated dynamic temporal properties of transitions between hidden brain states. A powerful feature of HMM is that it provides moment-by-moment estimates of the probability of either switching between latent states or staying within the same brain state. We computed the state transition matrix for each participant and first examined the likelihood that a brain state at time instance t remained within the same brain state in the previous time t − 1. This analysis revealed that all 10 brain states are *sticky*: i.e., states are not volatile from one-time step to another. (Fig. 4 C). These findings are important because they suggest that the latent brain states are stable over time. We also include in Fig. 4D the state transition probability without considering the self-transition (i.e., factoring out sticky states of the same kind, to view other possible relationships between states).

### Brain states and their interpretations

States were defined with HMM via network selected modularity extracted from each of the 4 networks included this study. Consistent with past research (Anderson, et al., 2014) we operationalized several reliable patterns of neural activity present in the HMM models. HMM is an exploratory approach, and requires choosing an artificial number of states beforehand; it is impossible to know *a priori* what the ideal state number will be, and often a portion of the resulting states may lack empirical relevance or be difficult to interpret clearly. This necessitates choosing a criteria for determining which states to focus on further (e.g., Liu, Amey, & Forbes, 2018; Vidaurre et al., 2018). We therefore applied two criteria in selecting relevant states: (1) network dominance or (2) high inter-network synchrony (which could, alternatively, reflect either a hybrid functional process or more efficient accommodation between two networks.

Four out of ten states were easily defined for their unique dominant network selected modularity value (≥50%) within each of the 4 networks we are interested in (see figure 5). For other states, their meanings are difficult to interpret, because they represented certain level of network activity for combination of 4 networks without revealing one dominant. To avoid the ambiguities and to make the analysis clearer, we only focused on these four well-defined states and their states properties, we call them SDMN, SFP, SEM and SER respectively. Figure 5A represents the 10 extracted states in terms of to what extent (in percentage) the selected modularity of each network contributes to the total variance of selected-modularity within each state.

**Figure 5.**
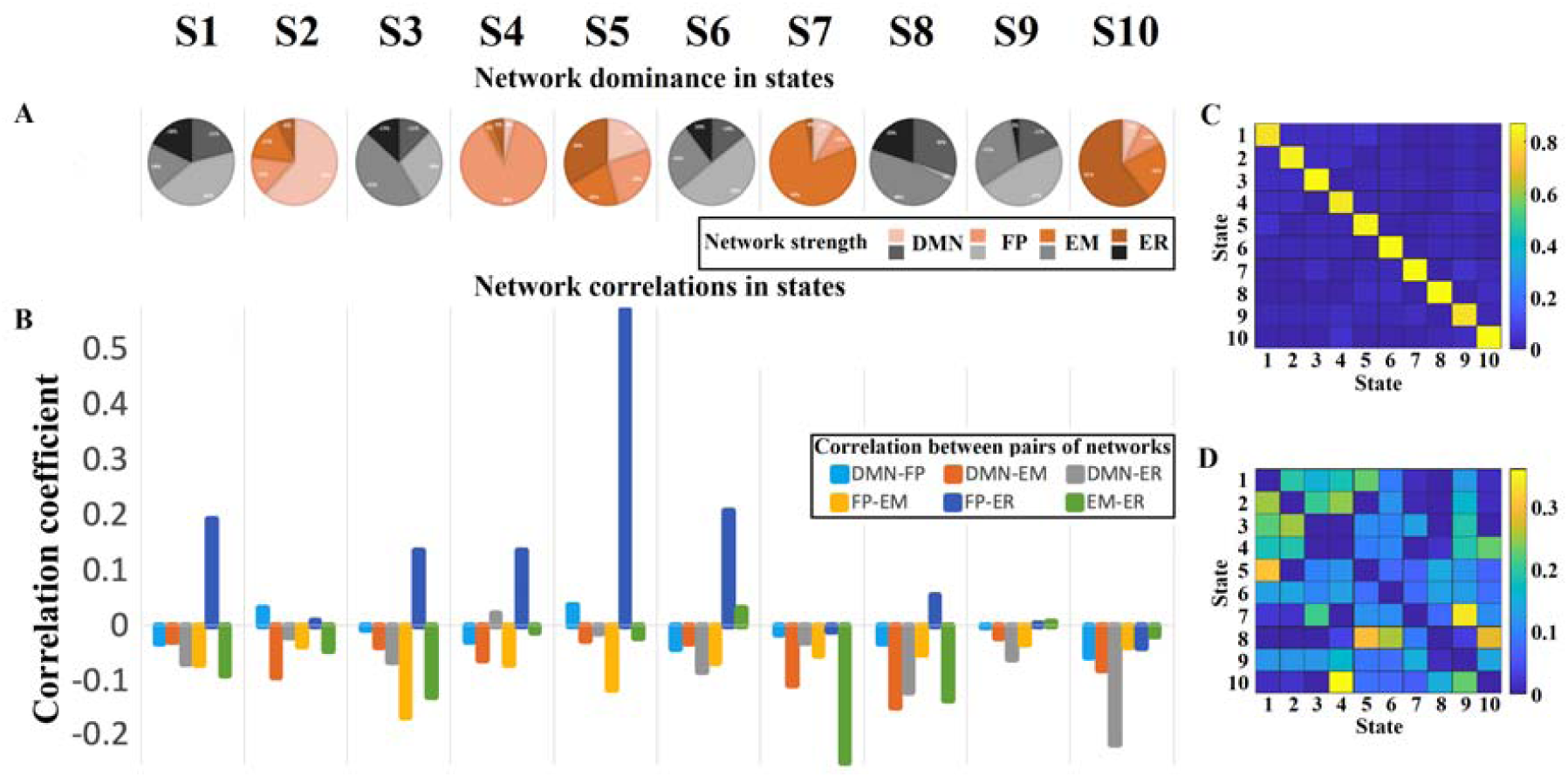
Latent brain states during math solving, their dynamic properties. A) Pie chart represents to what extent (in percentage) the selected modularity of each network contributes to the total variance of selected modularity within each state. B) Bar plot represents correlation between all pairs of network dynamics with in each of latent brain state. C) State transition matrix for latent brain states at time instance t remained within the same brain state in the previous time t − 1. D). State transition probability without considering the self-transition.

To gain further insight in to the role of states, we also investigated the network global synchrony by measuring whether oscillatory activity in each network varied in conjunction with one another or oscillated more randomly. We indexed synchrony by the Pearson’s correlation coefficient between all possible pairwise networks in a specific state (Figure 5B). Findings indicated that interestingly, in state 5, the frontoparietal and emotion regulation networks were strongly correlated (r = 0.57; corrected p < 0.0001; zscore = 4.01). Thus, we considered that state 5 might also play important role in solving math problems and hence state 5, named SFP-ER, analyzed together with other 4 well defined states. Other large synchrony between networks were found that in SEM state, EM network and ER network are highly anti-correlated (c = −0.31; corrected p < 0.0001; zscore = 3.17), indicating in SEM state the ER function was completely muted. Another finding is in SER state ER network and DM network are highly anti-correlated (c = −0.22; corrected p < 0.0001; zscore = 2.68). We think other states might be a different transition stage between the 4 functional states.

### Proportional occupancy of states predicts math performance

To probe the relation between latent brain states and task performance, we examined whether time-varying brain state changes could predict math performance. Meanwhile, instead of examining the predictive role of every single brain state, we set out to determine how all brain states, integrally, contribute to the success of math performance—i.e., relative ability for each of the states to predict performance. This is difficult to obtain by conventional linear or multiple linear models used in past studies (Taghia et al., 2018). Thus, in our study we predicted math performance using random forest regression (RFR), a supervised machine learning algorithm, which obtained a ranking of brain states importance by using the out-of-bag samples (Dvornek et al., 2018).

Specifically, RFR first calculated variables’ (brain states’) importance for predicting math performance using out-of-bag error (Figure 6). Higher values mean greater importance in prediction, while negative values indicate a given variable would harm the performance prediction (Bukhari, et al, 2016); said differently, the values provided can be conceived as the percentage by which a predictor contributes to the model. We then trained an RFR model to estimate math performance accuracy using occupancy rates of brain states that had importance scores over a 0.1 threshold, applied the model on unseen data to predict accuracy using 10-fold cross validation, and evaluated model performance by comparing estimated accuracy and observed scores across all the subjects. This analysis revealed a significant relation between predicted and actual accuracy in all participants (p < 0.0001, Pearson’s correlation, same below). Conducting the same analysis for each experimental cell of participants revealed significant relationships for men under control (p = 0.008); men under stress (p = 0.014), and women under stress (p = 0.011). No significant relation was found for women under control (r < 0).

**Figure 6.**
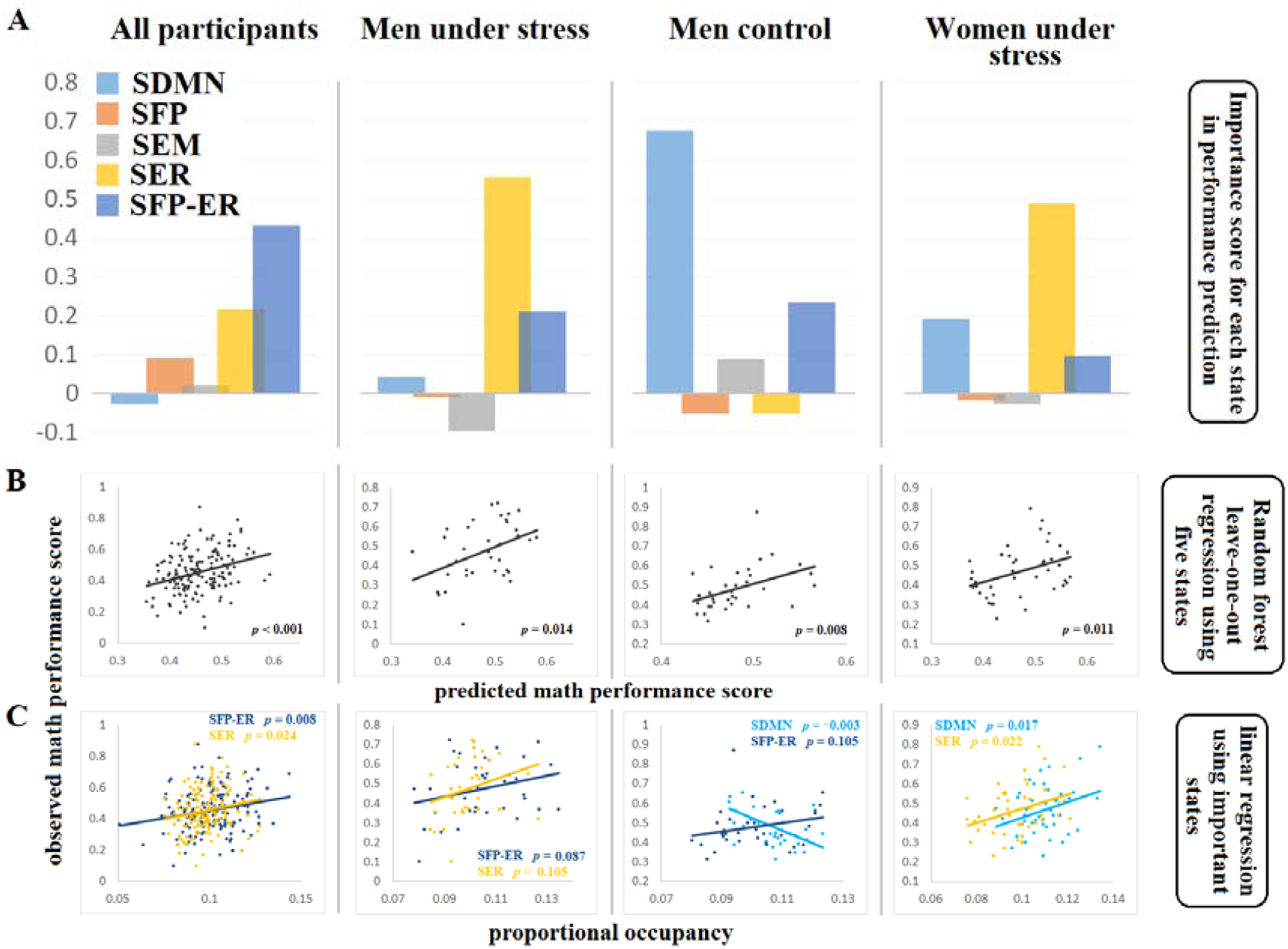
Proportional occupancy of states predicts math performance. A) Random forest analysis using out-of-bag samples calculated importance score of each brain states of interest in predicting math performance. B) RFR using important brain states can predict math performance score using 10-fold cross validation. C) The occupancy rate of individual brain states with higher importance score is associated with performance during math solving.

We then tested the hypothesis that the occupancy rates of individual brain states having higher importance scores would be associated with performance during math solving. We found that for all participants, math performance accuracy was positively correlated with the occupancy rate of FP-ER state (p = 0.008) and ER state (p = 0.024). For men under control, math performance accuracy was negatively correlated with the occupancy rate of the DMN state (p = 0.003), and marginally positively to the FP-ER state (p = 0.105). For men under stress, math performance accuracy was positively correlated with the occupancy rate of marginally for the ER state (p=0.087) and the FP-ER state (p = 0.107), For women under stress, math performance accuracy was positively correlated with the occupancy rate of the DMN state (p = 0.017) and the ER state (p=0.022)

### Mean lifetime of states predicts math performance

Next, we investigated the mean lifetime, another feature of the temporal evolution of latent brain states, in relation to math performance using the same analytic procedures described above. This analysis revealed a significant relationship between predicted and actual accuracy in all participants (p = 0.0004). Group-wise analysis revealed a significant relationship in men under control (p = 0.01) and in men under stress (p = 0.023). No significant relationships were found in the two women’s groups (*p*’s > 0.251). Predictor importance analysis indicated that for all participants, FP and EM states had importance scores above threshold. Group-wise analysis revealed that the DMN state played an important role in men under control, while the EM state was an important predictor for men under stress. Simple linear regression also suggested that for all participants, mean life of the FP state was positively associated with math performance (marginally significant: p = 0.082) and that the EM state was negatively associated with math performance (marginally significant: p = 0.118). For men under control, the DMN state negatively correlated with math performance (p < 0.001). For men under stress, the EM state negatively correlated with math performance (marginally significant: p = 0.103).

### Transition between different states

State transition probability is also an important index for describing the state propogation during the math solving process. It labels the temporally sequential relationship bet ween different states. In this study, the transition between two functional states (among the 5 states are interested in) may be concatenated with one or more transition states (among the 6 states). Thus, instead of measuring the direct transition between two functional states, we measured the average interval time between two functional states. We compared the 5 x 5 transition intervals between each pair of the functional states. Results revealed a main effect for gender (Men: 22.580.66 vs Women 20.260.63), corrected p = 0.027 and for condition (Control: 22.550.65 vs stress: 20.290.64) corrected p =0.038; from EM state to DM state. This result indicated female and stressed participants showed less time when they switch their brain from EM state to DM state. Simple two-tailed t-test was further conducted between women under stress and other participants in state transition from EM to DM, results revealed a significant effect (p = 0.009). This result is a further evidence DM state might play a role of emotion buffer when individuals were placed in stressful situation.

### Reliability test conducted: number of states between 9-11

To examine whether the number of states can impact the patterns we found, we evaluate the findings by using number of states of 9 and 11 respectively, instead of using only 10 states in HMM model. same analytic procedures described above were conducted.

To certify that states we picked up from different models are the same states, we tested how many time window we believed belong to same state for two models were overlapped. For example, we first found a default network dominant state in model A and another default network dominant state in model B. Then we selected all time windows belonged to this state in Model A, and all time windows belonged to this state in Model B. Then we measured how many time windows among them are same time windows, and their proportions on all the time windows selected. This step was conducted on all pairs of models, final overlap proportion is the average of all pairs of models for each state. Results indicated that for default dominant state, 69.7% of time windows were overlapped average from all pairs of models; for emotion regulation dominant network, 66.3%; for stress buffer dominant, 69.3%; and for task positive dominant, 67.3%. We can conclude from these results that the state we selected across models that represent the same function were accurate. See supplementary results for the cross state validations.

## Discussion

Results indicate that, when the brain is tasked with competing processing demands during problem solving, it accommodates through brief neural states that facilitate better cooperation between networks. Further, distinct neural states predicted better performance under conditions varying in difficulty and stressfulness. In line with prior work, FPN-dominant states reliably predicted better performance overall on easier math problems. However, other factors appeared more important for performance when difficulty increased, and diverging patterns emerged in as a function of stress condition. We will focus on difficult problems data for the majority of this discussion. Overall, we observed that states involving ER and FP-ER cooperation predicted better performance for more difficult problems. Examining further, when identity threats were introduced, FP-ER states remained, while suppressed DMN became important for the control group (men). Contrastingly, for the stress group (women), ER and DMN states predicted better performance. Thus, it appears that, through brief neural states, two routes to accommodation emerged in response to conflicting processing demands, which we venture to interpret in the following paragraphs.

Returning to our “tug-of-war” analogy (Fig. 1) it would seem that, overall, participants who incorporate ER and ER-FP network-dominant states in relation to better performance under increased difficulty are able to do so without “maxing out” their shared executive resources. It may be that ER states, for these subjects, are managing an arousal accompanying increased difficulty that is not negative in nature, and that ER-FP states then reflect more efficient integration between the two demands of executive attention to problems and regulation of internal states. Indeed, classic literature shows that arousal can in fact enhance performance.

Trends depart somewhat from this overall theme as we examine what happened with the addition of an evaluative threat. While men who performed well under identity threats continued to show ER-FPN states and *down*-regulated DMN, women’s states ceased to show FPN but rather added DMN. Why might this be the case? From identity threat literature, we know that men do not report feeling stressed and may even perform better on difficult math problems due to “stereotype lift”, or primed competence when gender is made salient; likewise, our males also did not evince signs of stress. The women in the evaluative threat group, on the other hand, did report feeling negative arousal. This suggests a difference in the experience of arousal between the stress and control group: a “challenge” (positive) versus “threat” effect, which is known to track with better or poorer performance under conditions of evaluative threat (Blascovich, 2013).

In light of this dynamic and given that both DM and ER states predicted better performance for the stress group, it suggests that DMN played a unique positive role— perhaps either in coping with evaluative threats by providing autobiographical buffer, or by more effectively managing internally oriented cognitions—thereby assisting in the regulation of negative arousal. This would fit with prior suggestions that DMN may play a role in coping, and it extends work demonstrating that DMN at rest contributed to more favorable self-assessments following stressful performance (Forbes, Duran, Schmader, & Allen, 2014). The fact that FP-ER states were no longer prominent in this condition may also reflect poorer coordination between these two functional systems. Said differently, in the “tug of war” between problem solving and self-regulation, ER may have gained priority. In contrast to the stress group, the men under evaluative threat conditions performed better in relation to states that *suppressed* DMN; while they retained the FP-ER states, ER states were no longer prominent. If the “tug of war” becomes more strained under evaluative threats, it follows that efficient cooperation and avoidance of anything that might tip the balance would be most critical in such situations.

It is important to add the caveat that the men in the evaluative threat condition may not have been truly be stress-free. but instead could have reflected lesser *levels* of stress compared to the women, which leant to a different response. This consideration stems from the fact that men exposed to evaluative stressors *did* differ from men under control—it became more important to down-regulate DMN. On the other hand, women benefitted from DMN-active states. Taken together, this may indicate that under evaluative threats, the brain seeks to mitigate intrusions when possible, but if necessary can opt for a “plan B”, that involves more of DM and ER. Putting things into perspective, women who displayed the DMN-dominant state fared *better* under stress, though performed worse overall than their male counterparts.

Finding that the brain is able to simultaneously accommodate demands for emotion regulation and executive attention to facilitate problem-solving performance lends new perspective to previous work, which largely documented a conflicting relationship. How can we reconcile these two stories? First, the present paradigm allowed us to deliberately manipulate stress. Therefore, the neural patterns we find in experimental cells are naturally characteristic of what predicts better performance *given* those contexts. This effectively allows us to tell a more complex story about problem difficulty and added stress, where before processes related to arousal and evaluative threat would have only surfaced as an individual difference measure in the general population. Thus, it makes sense here why some of these markers could actually be taken as *beneficial*, instead of muddying. This certainly does not detract from the fact that FPN processes remain the dominant ingredient for success on problem solving tasks. Rather, our aim was to spotlight what makes the difference under varying contexts with conflicting demands, all else considered.

Second, it is beneficial to briefly review what our data approach affords us (and it’s limitations). Applying HMM to measures of EEG network synchrony allows us a data-driven way to identify extremely fleeting neural states (ranging from, on average, 0.3-0.5 seconds), which may occur at any point in the time series. Hence, while the FPN dominant states evident for easy problem solving appear to share common ground with prior work, we are limited in our ability to make direct comparisons to prior work regarding of ERN and DMN findings. With respect to novel markers that emerged under increased difficulty and stress, we suggest these appear because of additional demands in those circumstances. Further, we warrant a measure of caution when interpreting EEG activation as regionally specific. We therefore minimize these drawbacks by using a carefully placed, high-density electrode array and adopting advanced Bayesian source localization (incorporating dSM operators) and confining analyses to cortical surface areas. Future research should complement these findings by applying spatially attuned, fMRI imaging—particularly given the importance of amygdala and medial temporal regions in arousal and self-referential processes (Forbes *et al.* 2018).

## Conclusion

Taken together, the findings present an intriguing window into how the brain might mitigate conflicts in processing demands that are often thought of as zero-sum, suggesting individual difference variables promoting success. By deliberately manipulating novel contextual variables of established problem-solving paradigms, we also move closer to more sophisticated, ecologically valid models of cognition. The resulting neural states we have reviewed underscore that network behavior may be highly nuanced at more micro time scales, involving the emergence of counterintuitive functional themes and inter-network cooperation. This emergent avenue of cognitive neuroscience offers a promising area for future research, with a variety of basic science applications.

## Supplemental Information

### Stress manipulation check: Stress and math performance score

To examine whether there were overall performance differences between conditions on math score performance, an initial 2 (Gender: Men or Women) x 2 (Condition: DMT or PST) factorial ANOVA was conducted on participants’ accuracy on the math feedback task (number correct/number attempted). This analysis yielded a main effect for gender, *F* (1, 156) =16.56, *p*<.001, *d*=.63. There were no other main effects or interaction (*p*’s>.38). Given the well documented effects of stereotype threat on performance (for a review see Schmader et al., 2008), however, planned contrasts were also conducted to compare DMT women’s performance on the math feedback task to the other three conditions. These analyses indicated that DMT women (i.e., those experiencing stereotype threat) performed worse on the math feedback task compared to the other three conditions, *t* (1, 156)=3.17, *p*=.002, *d*=.51.

With respect to the role of task difficulty, given the nature of our math feedback task, which was lengthy and contained easy, medium and difficult questions, it’s possible that these various problem types had variable effects on performance across groups. To examine this, a 2 (Gender: Men or Women) x 2 (Condition: DMT or PST) x 3 (Problem Type: easy, medium or difficult) mixed factors ANOVA with repeated measures on the latter variable was conducted. This analysis yielded a main effect for gender, *F*(1, 156)=12.38, *p*=.001, *d*=.55, that was qualified by a problem type by condition interaction, *F*(1, 156)=4.44, *p*=.01, *d*=.35, and problem type by gender interaction, *F*(1, 156)=3.41, *p*=.03, *d*=.29. Simple effects analyses using a Dunn-Sidak adjustment to control for multiple comparisons indicated that DMT women performed worse on easy, F(1, 156)=4.17, p=.04, d=.33, and difficult problems, F(1, 156)=4.73, p=.03, d=.35, compared to PST women. Men did not differ from one another with respect to condition, p’s>.23. DMT women also performed worse on easy, F(1, 156)=11.99, p=.001, d=.56, and moderately difficult problems, F(1, 156)=3.77, p=.05, d=.31, compared to DMT men. Women in the PST condition performed worse on easy, F(1, 156)=6.51, p=.01, d=.41, and moderate problems, F(1, 156)=5.64, p=.02, d=.38, compared to men in the PST condition. Interestingly only DMT women did not perform differently on easy, medium and difficult problem types, presumably because they underperformed across problem types in general (p’s>.12). All other groups showed the expected patterns, i.e., performing better on easy compared to moderate and difficult problems, and moderate compared to difficult problems (p’s<.04). Results overall provide supporting evidence that the stereotype threat manipulation was successful.

### FPN in easy problem predict performance

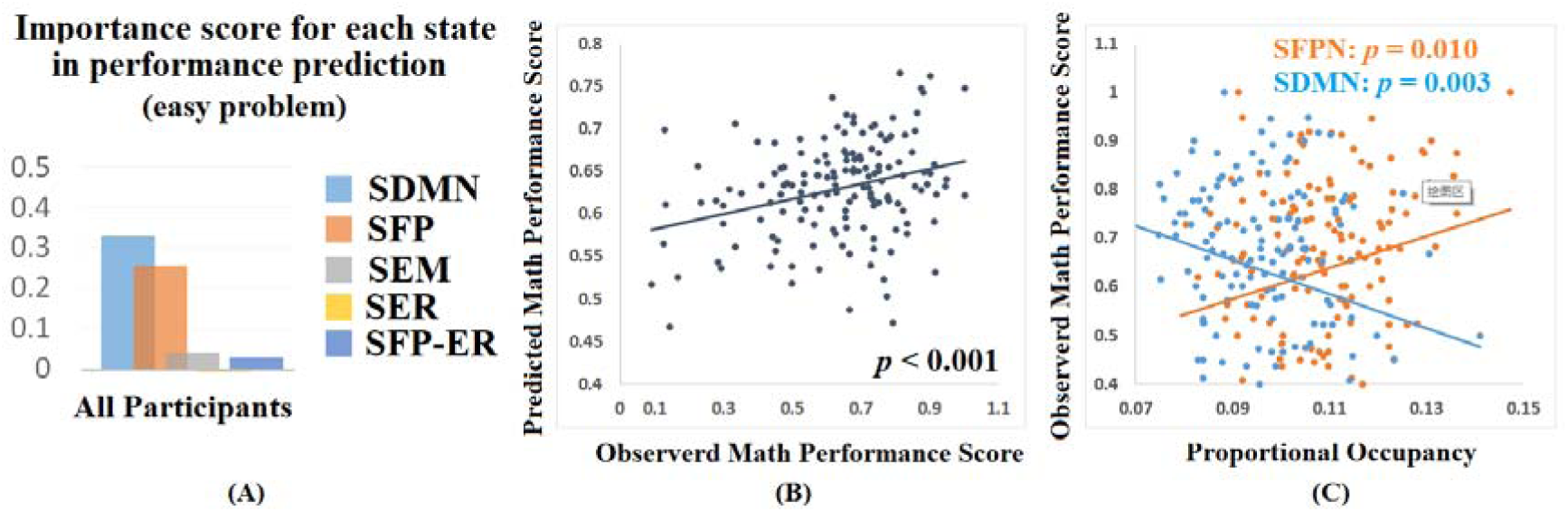

### Reliable test was conducted on number of states between 9-11

**State analysis for 11 states**

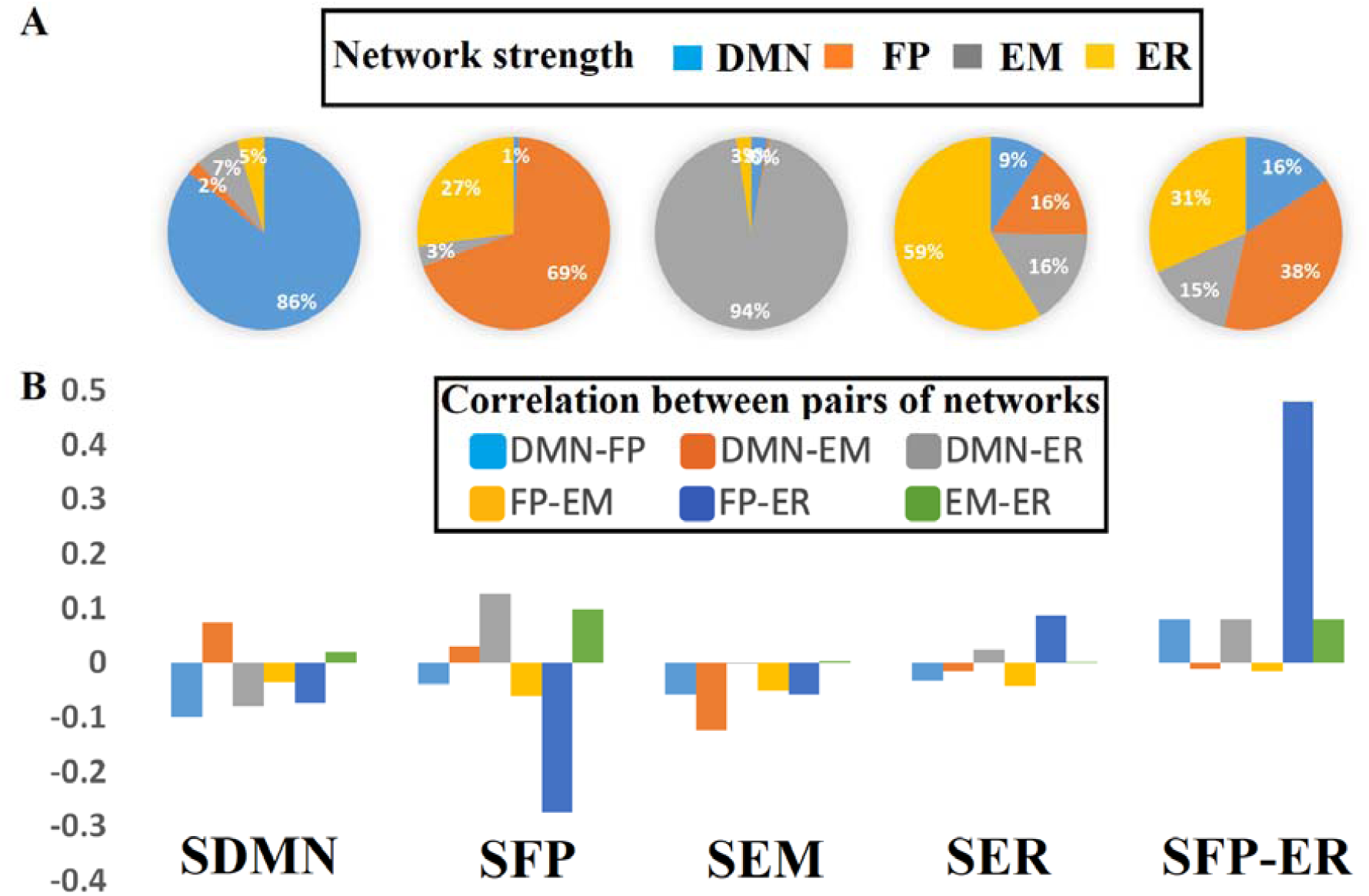

**2, Same figures for other states.**

**3, explanation of the meaning for all other states.**

### Proportional occupancy of brain states in different conditions

A 2 (condition: stress or control) x 2 (gender: male or female) between factors ANOVA was conducted on proportional occupancy of the four brain states we are interested in. There was also a two-way interaction between condition and gender for EM state (corrected p = 0.063), Simple effect analyses indicated that stressed women showed more EM state (M=7,7%, std=1.8%) in comparison to stressed men (M=7.2%, std=1.6%), p = 0.009. Simple effects were not found in control between men and women. No other main or effects were significant (corrected p’s > 0.91). No other main or interaction effects were significant for other states (corrected p > 0.117).

### Proportional occupancy of states predicts math performance

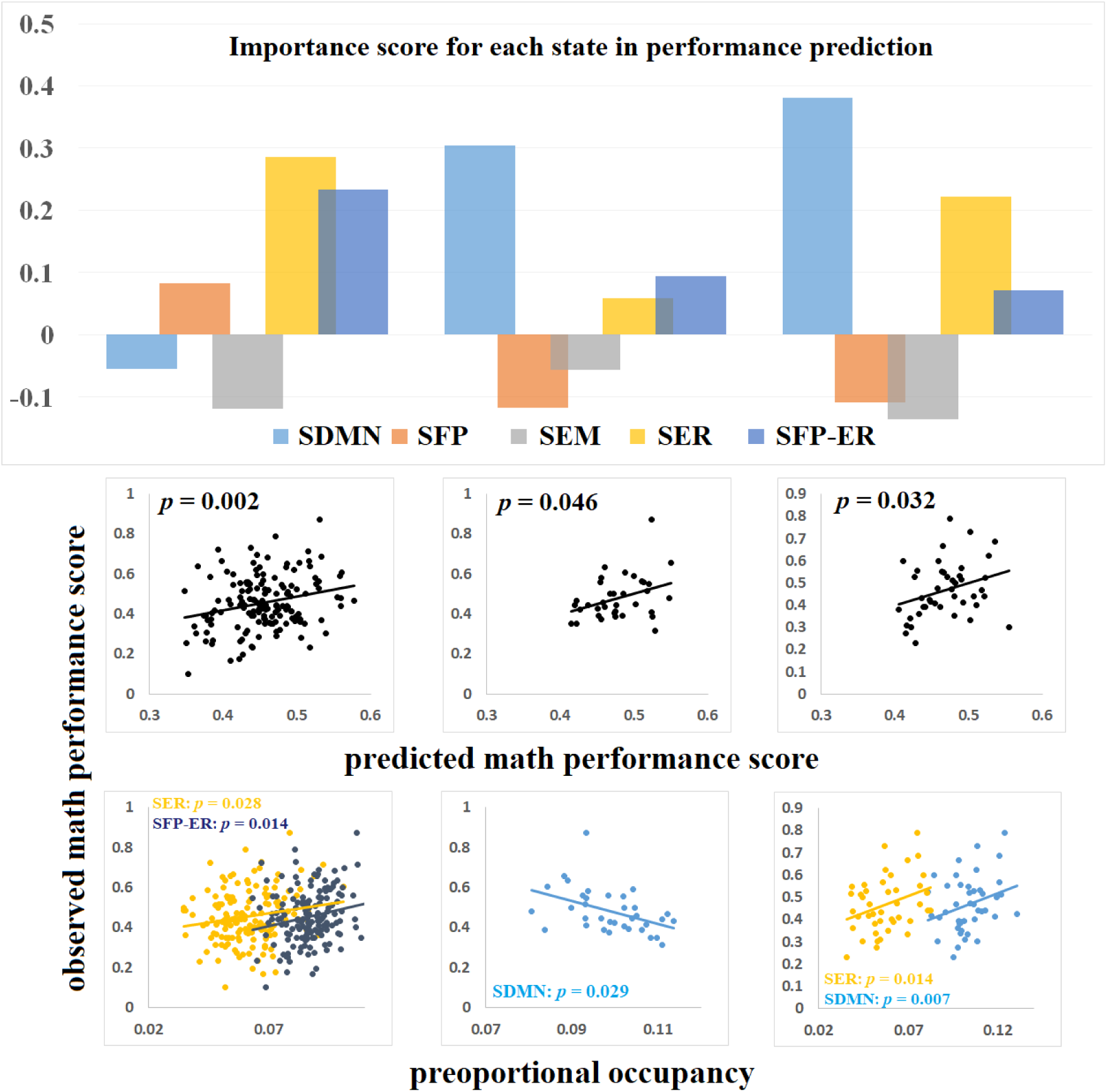

This analysis revealed a significant relation between predicted and actual accuracy for DM state (in negative relationship, corrected p = 0.041, Pearson’s correlation) for men under control. Thus, the dominant DM state is a behaviorally harmful brain state for math solving performance—the more time spent in this brain state the worse math task performance in people without any stress context.

This analysis also revealed a marginal significant relation between predicted and actual accuracy for DM state (in positive relationship, corrected p = 0.043, Pearson’s correlation) and ER state (in negative relationship, corrected p = 0.271, Pearson’s correlation) for women under stress. Default network has been found in past that buffers stress (Brewer et al., 2011), emotion regulation network has straightforward effect on stress regulation. Thus, we hypotheses these two states both plays a stress buffer function during math solving process, hence the combination of the two states would help suppress the stress emotion generated and in turn increase the math performance for individuals in stressful context. To investigate this, a new state was created by combining the two states mentioned above, and was named as stress buffer state (SB). Same analysis was conducted on the new stress buffer state and math solving accuracy. Significant effects were found were found for women under stress (p = 0.026). Thus, the dominant stress buffer state is a behaviorally optimal brain state for math performance for people in stressful context — the more time spent in this brain state the better math task performance.

### Mean interval of brain states in different conditions

A 2 (condition: stress or control) x 2 (gender: male or female) between factors ANOVA was conducted on mean interval time of the four brain states we are interested in. There was also a two-way interaction between condition and gender for EM state (corrected p = 0.056), Simple effect analyses indicated that stressed women showed less interval time between two EM state (M=193.9ms, std=6.4ms) in comparison to stressed men (M=214.8.8ms, std=8.1ms), p = 0.006. Simple effects were not found in control between men and women. No other main or effects were significant (corrected p’s > 0.23). There was also a difference between condition for ER state (corrected p = 0.063), Simple effect analyses indicated that stressed participants showed less interval time between two ER state (M=187.7ms, std=6.4ms) in comparison to control participants (M=205.4ms, std=9.2ms). No other effects were found (p > 0.34). No other effects were found in other states(p>0.33).

### Mean lifetime predicts math performance

Next, we investigated the mean lifetime, another key feature of temporal evolution of latent brain states, in relation to math performance using the same analytic procedures described above. We found that the mean lifetimes of DM brain states in the men under control negatively predicted math performance accuracy (corrected p = 0.109). Mean lifetime of the DM state was also a most robust predictor of the underperformance in the math task.

### Transition between different states

State transition probability is also an important index for describing the state propogation during the math solving process. It labels the temporally sequential relationship bet ween different states. In this study, the transition between two functional states (among the 4 states are interested in) may be concatenated with one or more transition states (among the 6 states). Thus, instead of measuring the direct transition between two functional states, we measured the average interval time between two functional states. We compared the 4 x 4 transition intervals between each pair of the functional states. Results revealed a main effect for gender (Men: 24.51+0.53 vs Women 22.34+0.49), corrected p = 0.032 and for condition (Control: 23.66+0.50 vs stress: 22.18+ 0.58) corrected p =0.058; from EM state to DM state. This result indicated female and stressed participants showed less time when they switch their brain from EM state to DM state. Simple two-tailed t-test was further conducted between women under stress and other participants in state transition from EM to DM, results revealed a significant effect (p = 0.021). This result is a further evidence DM state might play a role of emotion buffer when individuals were placed in stressful situation.

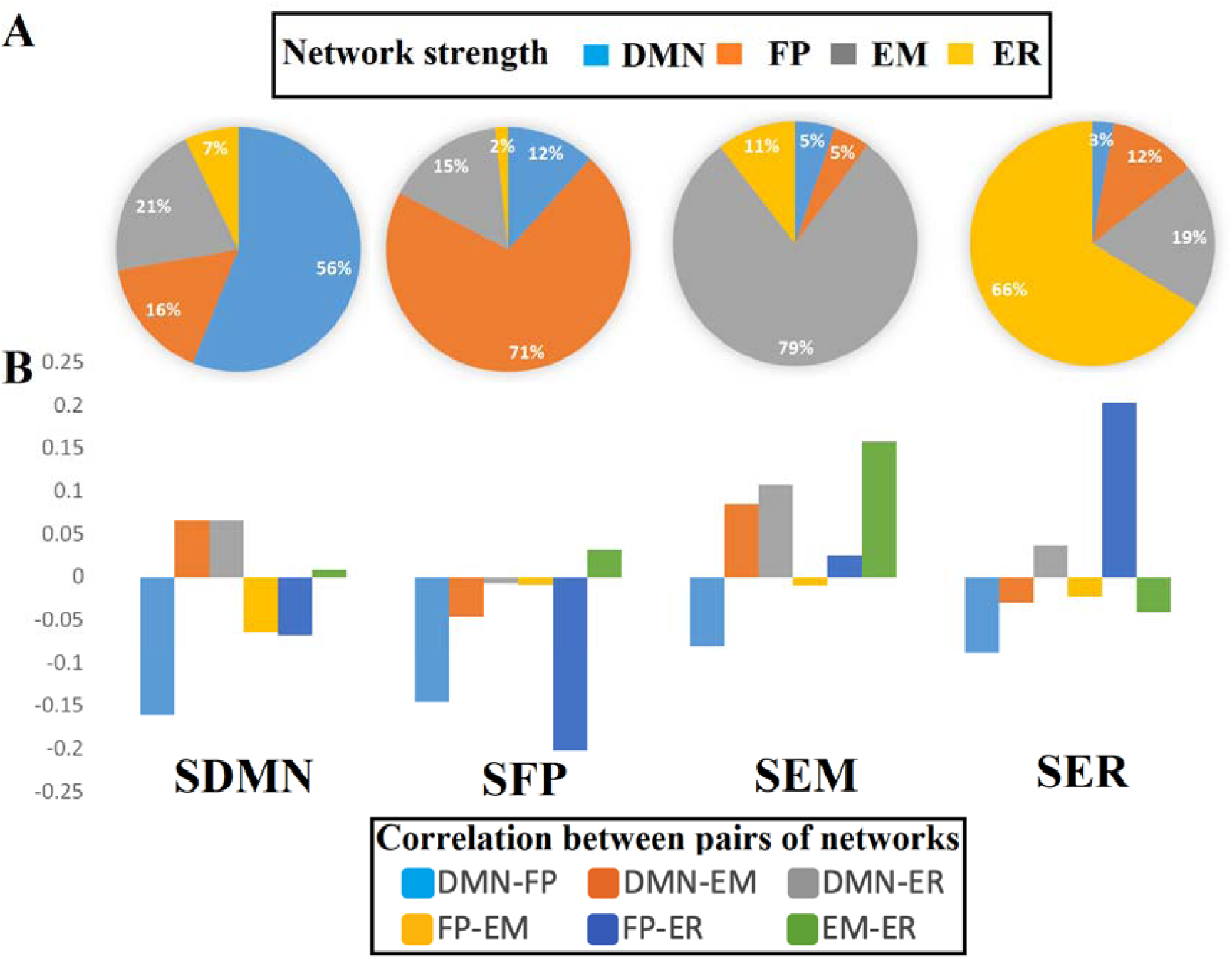

### Proportional occupancy of brain states in different conditions

A 2 (condition: stress or control) x 2 (gender: male or female) between factors ANOVA was conducted on proportional occupancy of the four brain states we are interested in. There was also a two-way interaction between condition and gender for EM state (corrected p = 0.043), Simple effect analyses indicated that stressed women showed more EM state (M=9.8%, std=2.1%) in comparison to stressed men (M=9.2%, std=2.2%), p = 0.009. Simple effects were not found in control between men and women. No other main or effects were significant (corrected p’s > 0.54). No other main or interaction effects were significant for other states (corrected p > 0.224).

### Proportional occupancy of states predict math performance

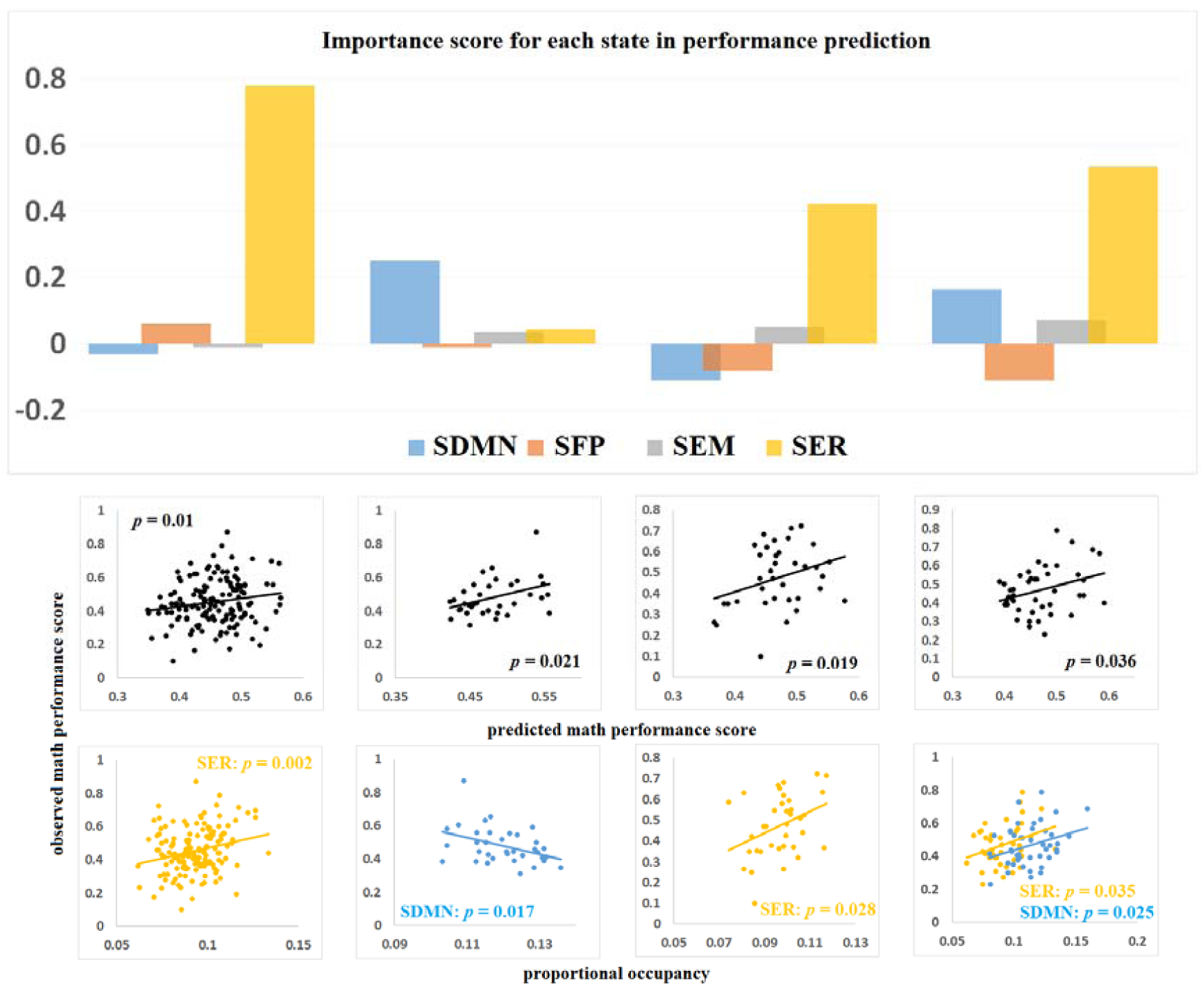

This analysis revealed a significant relation between predicted and actual accuracy for ER state (in negative relationship, corrected p = 0.075, Pearson’s correlation) for men under control. Thus, the dominant DM state is a behaviorally harmful brain state for math solving performance—the more time spent in this brain state the worse math task performance in people without any stress context.

This analysis also revealed a marginal significant relation between predicted and actual accuracy for DM state (in positive relationship, corrected p = 0.65, Pearson’s correlation) and ER state (in negative relationship, corrected p = 0.035, Pearson’s correlation) for women under stress. Default network has been found in past that buffers stress (Brewer et al., 2011), emotion regulation network has straightforward effect on stress regulation. Thus, we hypotheses these two states both plays a stress buffer function during math solving process, hence the combination of the two states would help suppress the stress emotion generated and in turn increase the math performance for individuals in stressful context. To investigate this, a new state was created by combining the two states mentioned above, and was named as stress buffer state (SB). Same analysis was conducted on the new stress buffer state and math solving accuracy. Significant effects were found were found for women under stress (p = 0.013). Thus, the dominant stress buffer state is a behaviorally optimal brain state for math performance for people in stressful context — the more time spent in this brain state the better math task performance.

### Mean interval of brain states in different conditions

A 2 (condition: stress or control) x 2 (gender: male or female) between factors ANOVA was conducted on mean interval time of the four brain states we are interested in. There was also a two-way interaction between condition and gender for EM state (corrected p = 0.115), Simple effect analyses indicated that stressed women showed less interval time between two EM state (M=46.21ms, std=1.38ms) in comparison to stressed men (M=50.03ms, std=1.51ms), p = 0.002. And compared to women under control (M=50.251,std=1.46), p=0.001. Simple effects were not found in control between men and women. No other main or effects were significant (corrected p’s > 0.23). There was also a two-way interaction between condition and gender for ER state (corrected p = 0.003), Simple effect analyses indicated that stressed women showed less interval time between two ER state (M=50.68ms, std=1.44ms) in comparison to control women (M=56.9ms, std=1.53ms). No other effects were found (p > 0.44). No other effects were found in other states(p>0.37).

### Mean lifetime predict math performance

Next, we investigated the mean lifetime, another key feature of temporal evolution of latent brain states, in relation to math performance using the same analytic procedures described above. We found that the mean lifetimes of DM brain states in the men under control negatively predicted math performance accuracy (corrected p = 0.074). Mean lifetime of the DM state was also a most robust predictor of the underperformance in the math task.

### Transition between different states

No significant effects were found for any of the states in any of the groups.

## Notes

### Competing Interest Statement

The authors have declared no competing interest.

## Reference

Aftanas, L., & Golosheykin, S. (2005). Impact of regular meditation practice on EEG activity at rest and during evoked negative emotions. International Journal of Neuroscience, 115(6), 893–909.

Allen, E. A., Damaraju, E., Plis, S. M., Erhardt, E. B., Eichele, T., & Calhoun, V. D. (2014). Tracking whole-brain connectivity dynamics in the resting state. Cerebral cortex, 24(3), 663–676.

Amey, R., Leitner, J., Liu, M., & Forbes, C. (2018). Neural Mechanisms Associated with Semantic and Basic Self-Oriented Memory Processes Interact to Modulate Self-Esteem. bioRxiv, 350926.

Anderson, J. R., Lee, H. S., & Fincham, J. M. (2014). Discovering the structure of mathematical problem solving. NeuroImage, 97, 163–177.

Bassett, D. S., Wymbs, N. F., Porter, M. A., Mucha, P. J., Carlson, J. M., & Grafton, S. T. (2011). Dynamic reconfiguration of human brain networks during learning. Proceedings of the National Academy of Sciences, 108(18), 7641–7646.

Bertolero, M. A., Yeo, B. T., & D’Esposito, M. (2015). The modular and integrative functional architecture of the human brain. Proceedings of the National Academy of Sciences, 112(49), E6798–E6807.

Buhle, J. T., Silvers, J. A., Wager, T. D., Lopez, R., Onyemekwu, C., Kober, H., … & Ochsner, K. N. (2014). Cognitive reappraisal of emotion: a meta-analysis of human neuroimaging studies. Cerebral cortex, 24(11), 2981–2990.

Bukhari, Q., Borsook, D., Rudin, M., & Becerra, L. (2016). Random Forest Segregation of Drug Responses May Define Regions of Biological Significance. Frontiers in computational neuroscience, 10, 21.

Cadinu, M., Maass, A., Rosabianca, A., & Kiesner, J. (2005). Why do women underperform under stereotype threat? Evidence for the role of negative thinking. Psychological science, 16(7), 572–578.

Cavanagh, J. F., & Frank, M. J. (2014). Frontal theta as a mechanism for cognitive control. Trends in cognitive sciences, 18(8), 414–421.

Celeux, G. (2007). Mixture models for classification. In Advances in data analysis (pp. 3–14). Springer, Berlin, Heidelberg.

Chen, P., Liu, R., Li, Y., & Chen, L. (2016). Detecting critical state before phase transition of complex biological systems by hidden Markov model. Bioinformatics, 32(14), 2143–2150.

Chen, J. L., Ros, T., & Gruzelier, J. H. (2013). Dynamic changes of ICA-derived EEG functional connectivity in the resting state. Human brain mapping, 34(4), 852–868.

Cheng, H., Linhares, B. M., Yu, W., Cardenas, M. G., Ai, Y., Jiang, W., … & Cierpicki, T. (2018). Identification of thiourea-based inhibitors of the B-cell lymphoma 6 BTB domain via NMR-based fragment screening and computer-aided drug design. Journal of medicinal chemistry, 61(17), 7573–7588.

Dale, A. M., Fischl, B., & Sereno, M. I. (1999). Cortical surface-based analysis: I. Segmentation and surface reconstruction. Neuroimage, 9(2), 179–194

Dale, A. M., Liu, A. K., Fischl, B. R., Buckner, R. L., Belliveau, J. W., Lewine, J. D., & Halgren, E. (2000). Dynamic statistical parametric mapping: combining fMRI and MEG for high-resolution imaging of cortical activity. Neuron, 26(1), 55–67.

Desikan, R. S., Ségonne, F., Fischl, B., Quinn, B. T., Dickerson, B. C., Blacker, D., … & Albert, M. S. (2006). An automated labeling system for subdividing the human cerebral cortex on MRI scans into gyral based regions of interest. Neuroimage, 31(3), 968–980.

Dvornek, N. C., Yang, D., Venkataraman, A., Ventola, P., Staib, L. H., Pelphrey, K. A., & Duncan, J. S. (2018). Prediction of autism treatment response from baseline fmri using random forests and tree bagging. arXiv preprint 1805.09799.

Eldawud, R., Reitzig, M., Opitz, J., Rojansakul, Y., Jiang, W., Nangia, S., & Dinu, C. Z. (2016). Combinatorial approaches to evaluate nanodiamond uptake and induced cellular fate. Nanotechnology, 27(8), 085107.

Etkin, A., Büchel, C., & Gross, J. J. (2015). The neural bases of emotion regulation. Nature reviews neuroscience, 16(11), 693–700.

Ertl, M., Hildebrandt, M., Ourina, K., Leicht, G., & Mulert, C. (2013). Emotion regulation by cognitive reappraisal—the role of frontal theta oscillations. NeuroImage, 81, 412–421.

Ferri, F., Tajadura-Jiménez, A., Väljamäe, A., Vastano, R., & Costantini, M. (2015). Emotion-inducing approaching sounds shape the boundaries of multisensory peripersonal space. Neuropsychologia, 70, 468–475.

Gross, J. J. (1998). Antecedent-and response-focused emotion regulation: divergent consequences for experience, expression, and physiology. Journal of personality and social psychology, 74(1), 224.

Gross, J. J., & Feldman Barrett, L. (2011). Emotion generation and emotion regulation: One or two depends on your point of view. Emotion review, 3(1), 8–16.

Goldin, P. R., McRae, K., Ramel, W., & Gross, J. J. (2008). The neural bases of emotion regulation: reappraisal and suppression of negative emotion. Biological psychiatry, 63(6), 577–586.

Forbes, C. E., Amey, R., Magerman, A. B., Duran, K., & Liu, M. (2018). Stereotype-based stressors facilitate emotional memory neural network connectivity and encoding of negative information to degrade math self-perceptions among women. Social cognitive and affective neuroscience, 13(7), 719–740.

Forbes, C. E., Leitner, J. B., Duran-Jordan, K., Magerman, A. B., Schmader, T., & Allen, J. J. (2015). Spontaneous default mode network phase-locking moderates performance perceptions under stereotype threat. Social cognitive and affective neuroscience, 10(7), 994–1002.

Forbes, C. E., Duran, K. A., Leitner, J. B., & Magerman, A. (2015). Stereotype threatening contexts enhance encoding of negative feedback to engender underperformance and anxiety. Social Cognition, 33(6), 605–625.

Gramfort, A., Luessi, M., Larson, E., Engemann, D. A., Strohmeier, D., Brodbeck, C., … & Hämäläinen, M. (2013). MEG and EEG data analysis with MNE-Python. Frontiers in neuroscience, 7, 267.

Gramfort, A., Luessi, M., Larson, E., Engemann, D. A., Strohmeier, D., Brodbeck, C., … & Hämäläinen, M. S. (2014). MNE software for processing MEG and EEG data. Neuroimage, 86, 446–460.

Hutchison, R. M., Womelsdorf, T., Allen, E. A., Bandettini, P. A., Calhoun, V. D., Corbetta, M., … & Handwerker, D. A. (2013). Dynamic functional connectivity: promise, issues, and interpretations. Neuroimage, 80, 360–378.

Jiang, W., Luo, J., & Nangia, S. (2015). Multiscale approach to investigate self-assembly of telodendrimer based nanocarriers for anticancer drug delivery. Langmuir, 31(14), 4270–4280.

Jiang, W., Wang, X., Guo, D., Luo, J., & Nangia, S. (2016). Drug-specific design of telodendrimer architecture for effective doxorubicin encapsulation. The Journal of Physical Chemistry B, 120(36), 9766–9777.

Jordano, M. L., & Touron, D. R. (2017). Stereotype threat as a trigger of mind-wandering in older adults. Psychology and Aging, 32(3), 307.

Kohn, N., Eickhoff, S. B., Scheller, M., Laird, A. R., Fox, P. T., & Habel, U. (2014). Neural network of cognitive emotion regulation—an ALE meta-analysis and MACM analysis. Neuroimage, 87, 345–355.

LeDoux, J. E. (2000). Emotion circuits in the brain. Annual review of neuroscience, 23(1), 155–184.

Leitner, J. B., & Forbes, C. E. (2015). The role of implicit mechanisms in buffering self-esteem from social threats. In Exploring Implicit Cognition: Learning, Memory, and Social Cognitive Processes (pp. 183-204). IGI Global.

Lin, F. H., Belliveau, J. W., Dale, A. M., & Hämäläinen, M. S. (2006). Distributed current estimates using cortical orientation constraints. Human brain mapping, 27(1), 1–13.

Liu, M., Amey, R. C., & Forbes, C. E. (2017). On the role of situational stressors in the disruption of global neural network stability during problem solving. Journal of cognitive neuroscience, 29(12), 2037–2053.

Liu, M., Kuo, C. C., & Chiu, A. W. (2011, August). Statistical threshold for nonlinear granger causality in motor intention analysis. In 2011 Annual International Conference of the IEEE Engineering in Medicine and Biology Society (pp. 5036-5039). IEEE.

Liu, M. T., Kuo, C. C., & Chiu, A. W. (2013). Non-linear Granger causality and its frequency decomposition in decoding human upper limb movement intentions. International Journal of Biomedical Engineering and Technology 34, 12(1), 1–25.

Liu, M., Amey, R. C., Magerman, A., Scott, M., & Forbes, C. (2020). The role of startle fluctuation and non-response startle reflex in tracking amygdala dynamics. bioRxiv.

Liu, M., Baker R. A., Amey, R. C., & Forbes, C. E. (2020). Context matters: Situational stress impedes functional reorganization of intrinsic brain connectivity during problem solving. bioRxiv.

Liu, M., & Wang, X. (2017). Beyond the ERPs—Startle Response is Better Outlined by Whole Brain and Spectral EEG Features. Journal of Psychiatry and Brain Science, 2(3).

Lisman, J. (2010). Working memory: the importance of theta and gamma oscillations. Current Biology, 20(11), R490–R492.

Liston, C., McEwen, B. S., & Casey, B. J. (2009). Psychosocial stress reversibly disrupts prefrontal processing and attentional control. Proceedings of the National Academy of Sciences, 106(3), 912–917.

Mantena, V., Jiang, W., Li, J., & McKenzie, R. (2009, April). Prostate cancer biomarker identification using MALDI-MS data: initial results. In 2009 IEEE/NIH Life Science Systems and Applications Workshop (pp. 116-119). IEEE.

Meunier, D., Lambiotte, R., & Bullmore, E. T. (2010). Modular and hierarchically modular organization of brain networks. Frontiers in neuroscience, 4, 200.

Nolan, H., Whelan, R., & Reilly, R. B. (2010). FASTER: fully automated statistical thresholding for EEG artifact rejection. Journal of neuroscience methods, 192(1), 152–162.

Ochsner, K. N., Bunge, S. A., Gross, J. J., & Gabrieli, J. D. (2002). Rethinking feelings: an FMRI study of the cognitive regulation of emotion. Journal of cognitive neuroscience, 14(8), 1215–1229.

Ochsner, K. N., & Gross, J. J. (2005). The cognitive control of emotion. Trends in cognitive sciences, 9(5), 242–249.

Ochsner, K. N., Ray, R. D., Cooper, J. C., Robertson, E. R., Chopra, S., Gabrieli, J. D., & Gross, J. J. (2004). For better or for worse: neural systems supporting the cognitive down-and up-regulation of negative emotion. Neuroimage, 23(2), 483–499.

Ossandón, T., Jerbi, K., Vidal, J. R., Bayle, D. J., Henaff, M. A., Jung, J., … & Lachaux, J. P. (2011). Transient suppression of broadband gamma power in the default-mode network is correlated with task complexity and subject performance. Journal of Neuroscience, 31(41), 14521–14530.

Ou, J., Xie, L., Wang, P., Li, X., Zhu, D., Jiang, R., … & Liu, T. (2013, November). Modeling brain functional dynamics via hidden Markov models. In 2013 6th International IEEE/EMBS Conference on Neural Engineering (NER) (pp. 569–572). IEEE.

Paré, D., Collins, D. R., & Pelletier, J. G. (2002). Amygdala oscillations and the consolidation of emotional memories. Trends in cognitive sciences, 6(7), 306–314.

Phan, K. L., Taylor, S. F., Welsh, R. C., Decker, L. R., Noll, D. C., Nichols, T. E., … & Liberzon, I. (2003). Activation of the medial prefrontal cortex and extended amygdala by individual ratings of emotional arousal: a fMRI study. Biological psychiatry, 53(3), 211–215.

Phillips, M. L., Ladouceur, C. D., & Drevets, W. C. (2008). A neural model of voluntary and automatic emotion regulation: implications for understanding the pathophysiology and neurodevelopment of bipolar disorder. Molecular psychiatry, 13(9), 833–857.

Preti, M. G., Bolton, T. A., & Van De Ville, D. (2017). The dynamic functional connectome: State-of-the-art and perspectives. Neuroimage, 160, 41–54.

Rabiner, L. R. (1989). A tutorial on hidden Markov models and selected applications in speech recognition. Proceedings of the IEEE, 77(2), 257–286.

Raghavachari, S., Kahana, M. J., Rizzuto, D. S., Caplan, J. B., Kirschen, M. P., Bourgeois, B., … & Lisman, J. E. (2001). Gating of human theta oscillations by a working memory task. Journal of Neuroscience, 21(9), 3175–3183.

Rudie, J. D., Brown, J. A., Beck-Pancer, D., Hernandez, L. M., Dennis, E. L., Thompson, P. M., … & Dapretto, M. J. N. C. (2013). Altered functional and structural brain network organization in autism. NeuroImage: clinical, 2, 79–94.

Sauseng, P., Klimesch, W., Schabus, M., & Doppelmayr, M. (2005). Fronto-parietal EEG coherence in theta and upper alpha reflect central executive functions of working memory. International journal of Psychophysiology, 57(2), 97–103.

Schmader, T., Johns, M., & Forbes, C. (2008). An integrated process model of stereotype threat effects on performance. Psychological review, 115(2), 336.

Silvers, J. A., Wager, T. D., Weber, J., & Ochsner, K. N. (2015). The neural bases of uninstructed negative emotion modulation. Social cognitive and affective neuroscience, 10(1), 10–18.

Simon, H. A. (1991). The architecture of complexity. In Facets of systems science (pp. 457–476). Springer, Boston, MA.

Spreng, R. N., Mar, R. A., & Kim, A. S. (2009). The common neural basis of autobiographical memory, prospection, navigation, theory of mind, and the default mode: a quantitative metaanalysis. Journal of cognitive neuroscience, 21(3), 489–510.

Suk, H. I., Wee, C. Y., Lee, S. W., & Shen, D. (2016). State-space model with deep learning for functional dynamics estimation in resting-state fMRI. NeuroImage, 129, 292–307.

Taghia, J., Cai, W., Ryali, S., Kochalka, J., Nicholas, J., Chen, T., & Menon, V. (2018). Uncovering hidden brain state dynamics that regulate performance and decision-making during cognition. Nature communications, 9(1), 1–19.

Van Ast, V. A., Spicer, J., Smith, E. E., Schmer-Galunder, S., Liberzon, I., Abelson, J. L., & Wager, T. D. (2016). Brain mechanisms of social threat effects on working memory. Cerebral Cortex, 26(2), 544–556.

Vidaurre, D., Hunt, L. T., Quinn, A. J., Hunt, B. A., Brookes, M. J., Nobre, A. C., & Woolrich, M. W. (2018). Spontaneous cortical activity transiently organises into frequency specific phase-coupling networks. Nature communications, 9(1), 1–13.

Wager, T. D., Davidson, M. L., Hughes, B. L., Lindquist, M. A., & Ochsner, K. N. (2008). Prefrontal-subcortical pathways mediating successful emotion regulation. Neuron, 59(6), 1037–1050.

Yu, S. Z., & Kobayashi, H. (2003). An efficient forward-backward algorithm for an explicit-duration hidden Markov model. IEEE signal processing letters, 10(1), 11–14.

Yu, S. Z., & Kobayashi, H. (2006). Practical implementation of an efficient forward-backward algorithm for an explicit-duration hidden Markov model. IEEE Transactions on Signal Processing, 54(5), 1947–1951.

Yu, Q., Erhardt, E. B., Sui, J., Du, Y., He, H., Hjelm, D., … & Calhoun, V. D. (2015). Assessing dynamic brain graphs of time-varying connectivity in fMRI data: application to healthy controls and patients with schizophrenia. Neuroimage, 107, 345–355.

